# Bridge RNAs direct modular and programmable recombination of target and donor DNA

**DOI:** 10.1101/2024.01.24.577089

**Authors:** Matthew G. Durrant, Nicholas T. Perry, James J. Pai, Aditya R. Jangid, Januka S. Athukoralage, Masahiro Hiraizumi, John P. McSpedon, April Pawluk, Hiroshi Nishimasu, Silvana Konermann, Patrick D. Hsu

## Abstract

Genomic rearrangements, encompassing mutational changes in the genome such as insertions, deletions, or inversions, are essential for genetic diversity. These rearrangements are typically orchestrated by enzymes involved in fundamental DNA repair processes such as homologous recombination or in the transposition of foreign genetic material by viruses and mobile genetic elements (MGEs). We report that IS110 insertion sequences, a family of minimal and autonomous MGEs, express a structured non-coding RNA that binds specifically to their encoded recombinase. This bridge RNA contains two internal loops encoding nucleotide stretches that base-pair with the target DNA and donor DNA, which is the IS110 element itself. We demonstrate that the target-binding and donor-binding loops can be independently reprogrammed to direct sequence-specific recombination between two DNA molecules. This modularity enables DNA insertion into genomic target sites as well as programmable DNA excision and inversion. The IS110 bridge system expands the diversity of nucleic acid-guided systems beyond CRISPR and RNA interference, offering a unified mechanism for the three fundamental DNA rearrangements required for genome design.

## INTRODUCTION

Evolution has dedicated a vast number of enzymes to rearrange and diversify the genome. From the complex interplay of eukaryotic DNA repair pathways to the arms race between prokaryotes and their viruses, DNA manipulations enable the emergence and functional specialization of new genes, the development of immunity (Tonegawa, 1983), and the opportunistic spread of viruses and mobile genetic elements (MGEs) (McCLINTOCK, 1951). MGEs are abundant throughout all domains of life and often mobilize themselves via a transposase, integrase, homing endonuclease, or recombinase. These enzymes typically recognize DNA via protein-DNA contacts and can be broadly classified by their target sequence specificity, which ranges from site-specific (e.g. Cre and Bxb1 recombinases) (Hoess et al., 1982; Russell et al., 2006) to semi-random (e.g. Tn5 and Piggybac transposases) (Blot et al., 1993; Fraser et al., 1995).

Insertion sequence (IS) elements are among the most minimal autonomous MGEs. Abundantly found across bacteria and archaea, these compact DNA sequences typically encode only the components necessary for transposition. For many characterized IS elements, mobility is enabled by a self-encoded transposase that recognizes terminal inverted repeats (TIRs) through protein-DNA interactions (Siguier et al., 2015). IS elements have been categorized into approximately 28 families based on their transposition mechanisms, protein homology, and DNA sequence architecture, but they can be broadly grouped by the conserved catalytic residues of their encoded transposases. These include DDE, DEDD or HUH transposases, and, less frequently, serine or tyrosine transposases that are related to common phage integrases (Siguier et al., 2015).

IS110 family elements are cut-and-paste MGEs that scarlessly excise themselves from the genome and generate a circular form as part of their transposition mechanism (Higgins et al., 2007; Partridge & Hall, 2003). Given what is known about this mechanism and life-cycle, IS110 transposases are more accurately described as recombinases. While circular intermediates are found in other IS families, IS110 is the only one of these families that encodes the DEDD catalytic motif in their recombinase. Furthermore, the N-terminal DEDD domains of the IS110 recombinases share homology with RuvC Holliday junction resolvases, together suggesting a unique mechanism of action compared to other IS elements. IS110 elements typically lack TIRs and appear to integrate in a sequence-specific manner, often targeting repetitive elements in microbial genomes (Tobes & Pareja, 2006). While the mechanism of DNA recognition and recombination for IS110 elements remains unclear, previous studies have suggested that the non-coding ends of the element flanking the recombinase ORF may regulate recombinase expression (Müller et al., 2001; Perkins-Balding et al., 1999).

Here, we report that the IS110 circular form drives the expression of a non-coding RNA (ncRNA) with two distinct binding loops that separately recognize the IS110 DNA donor and its genomic integration target site. By bridging the donor and target DNA molecules through direct base-pairing interactions, the bispecific bridge RNA facilitates DNA recombination by the IS110 recombinase. Each binding loop of the bridge RNA can be independently reprogrammed to bind and recombine diverse DNA sequences. We further demonstrate that this modularity facilitates sequence-specific insertion, inversion, and excision. By using the first reprogrammable and single-effector RNA-guided enzyme function beyond simple nucleic-acid binding and cleavage, bridge recombination enables a generalizable mechanism for the 3 fundamental DNA rearrangements.

## RESULTS

### IS621 expresses a non-coding RNA that is bound by its recombinase

Given the intriguing but sparse information gathered from studies of individual IS110 elements, we sought to characterize IS110 elements through a systematic computational analysis of their sequence features. Previous reports have noted that IS110 elements encode ∼300-460 aa recombinases with an N-terminal DEDD RuvC-like domain (Pfam ID: PF01548) and a C-terminal domain with a highly conserved serine residue (Pfam ID: PF02371, **Fig. 1a**, **Supplementary Fig. 1a,b**) (Siguier et al., 2015; Tobiason et al., 2001). IS110 elements utilize a recombinase to scarlessly excise out of their genomic context, yielding a dsDNA circular form that is inserted into specific genomic target sequences such as repetitive extragenic palindromic (REP) elements, through an as yet unknown mechanism (**Fig. 1b**) (Choi et al., 2003; Higgins et al., 2007, 2009; Perkins-Balding et al., 1999). Recombination of the circular form and the target is centered around a short core sequence (∼1-5 nt long), which appears as a direct repeat immediately flanking the inserted element. The intervening sequences between the cores and the recombinase coding sequence (CDS) are defined as the left (LE) and right (RE) non-coding ends. A phylogenetic analysis of IS110 recombinases that we extracted from approximately 13 terabases of publicly available metagenomic and genomic sequence data demonstrates that they are highly diverse and widespread in both bacteria and archaea. It further highlights that only a small subset of IS110 elements have been cataloged by curated databases, and that even fewer have been described in detail in the literature (**Fig. 1c**).

**Figure 1:**
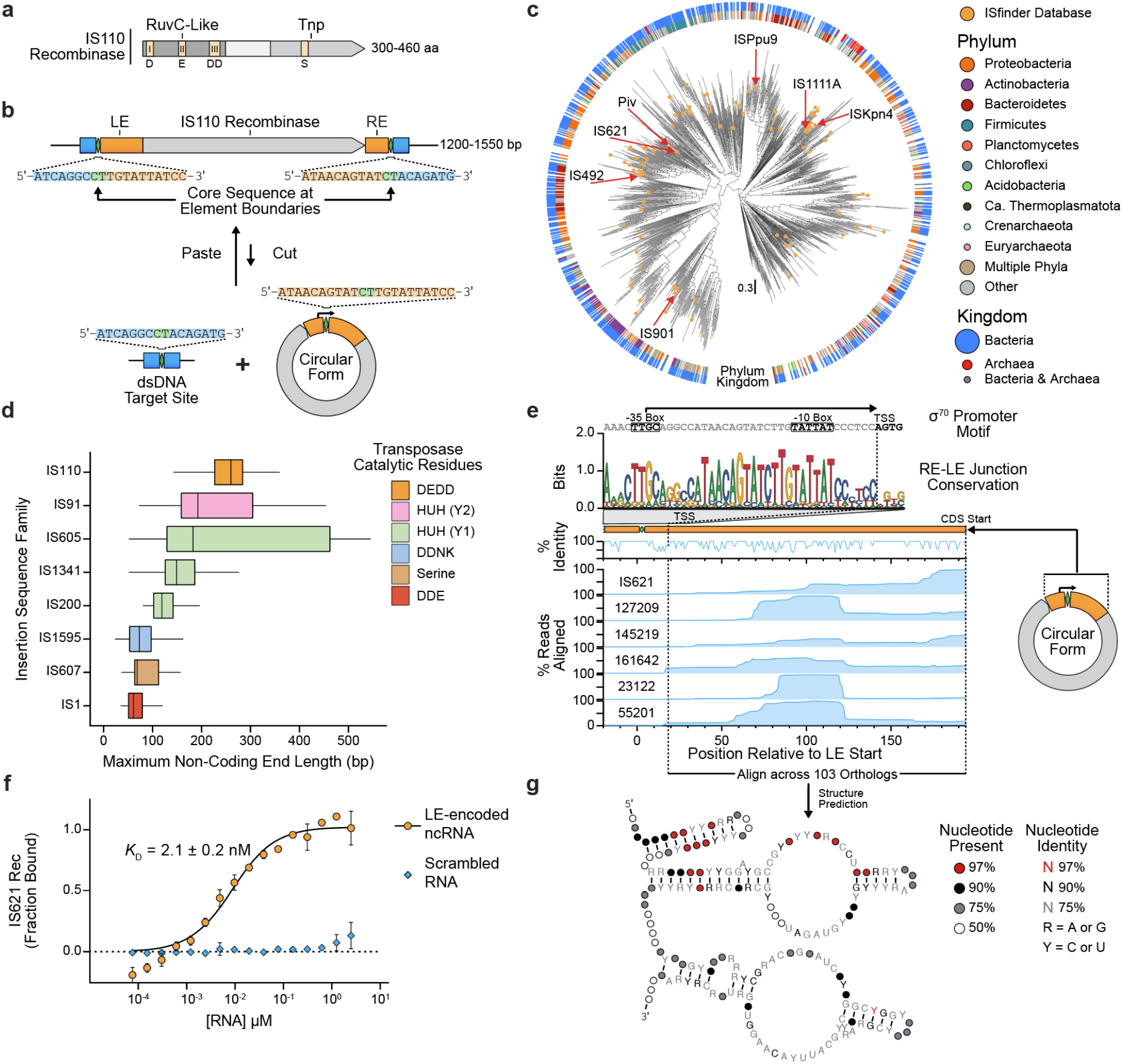
IS110 mobile genetic elements express a non-coding RNA that is bound by its encoded recombinase. (**a**) Schematic representation of the IS110 recombinase protein sequence. The RuvC-like and Tnp domains are highlighted along with the conserved catalytic residues. I, II, and III denote the three conserved and discontiguous subdomains of RuvC. (**b**) Schematic illustrating the structure and life-cycle of a typical IS110 element. Core sequences are depicted as green diamonds, the genomic target site is shown in blue and non-coding ends are orange. Sequences depicted are from IS621, an IS110 family member. (**c**) A midpoint-rooted phylogenetic tree constructed from 1,054 IS110 recombinase sequences. Tips indicated with a peach point denote IS110 family members cataloged in the ISfinder database (Siguier et al., 2006). The rings around the tree indicate the predominant taxonomic kingdom and phylum of each ortholog. The phylum and kingdom circles are sized according to the relative number of orthologs belonging to each division. (**d**) Distribution of non-coding end lengths across eight IS families with five different catalytic motifs. The maximum of the LE and RE lengths is plotted for each family. (**e**) Small RNA-seq coverage plot of the concatenated non-coding ends of IS621 and 5 related orthologs expressed from their endogenous promoter in *E. coli* (right). The top panel shows a sequence logo of the conservation of the 𝛔^70^ promoter motif. Percent nucleotide identity among the 6 orthologs is indicated across the top of the coverage plot. The vertical dotted line indicates the predicted transcription start site (TSS). (**f**) Microscale thermophoresis (MST) of a fluorescently labeled IS621 recombinase mixed with either WT or scrambled ncRNA to assess the equilibrium dissociation constant (*KD*). Mean ± SD of three technical replicates are shown. (**g**) Consensus secondary structure of ncRNAs constructed from 103 IS110 LE sequences. The predicted structure comprises a 5′ stem-loop and two large internal loops.

Analyzing the distribution of non-coding end lengths among IS110s and other IS families, we noted that IS110s have the longest median non-coding ends with a relatively narrow distribution, suggesting this feature is conserved across IS110 elements (**Fig. 1d**, **Supplementary Fig. 2a**). Upon excision, the circular form of the element reconstitutes a promoter across the core sequence of the concatenated RE-LE far upstream of the recombinase CDS (Müller et al., 2001; Perkins-Balding et al., 1999) (**Fig. 1b**), suggesting the possibility that a ncRNA could be expressed from this region. Previous reports have demonstrated that the non-coding ends of some other IS element families are transcribed into RNAs resembling CRISPR RNAs to guide endonuclease activity (Altae-Tran et al., 2021; Karvelis et al., 2021), and small RNAs have been thought to modulate recombinase expression for the IS110 family member ISPpu9 (Gómez-García et al., 2021).

**Figure 2:**
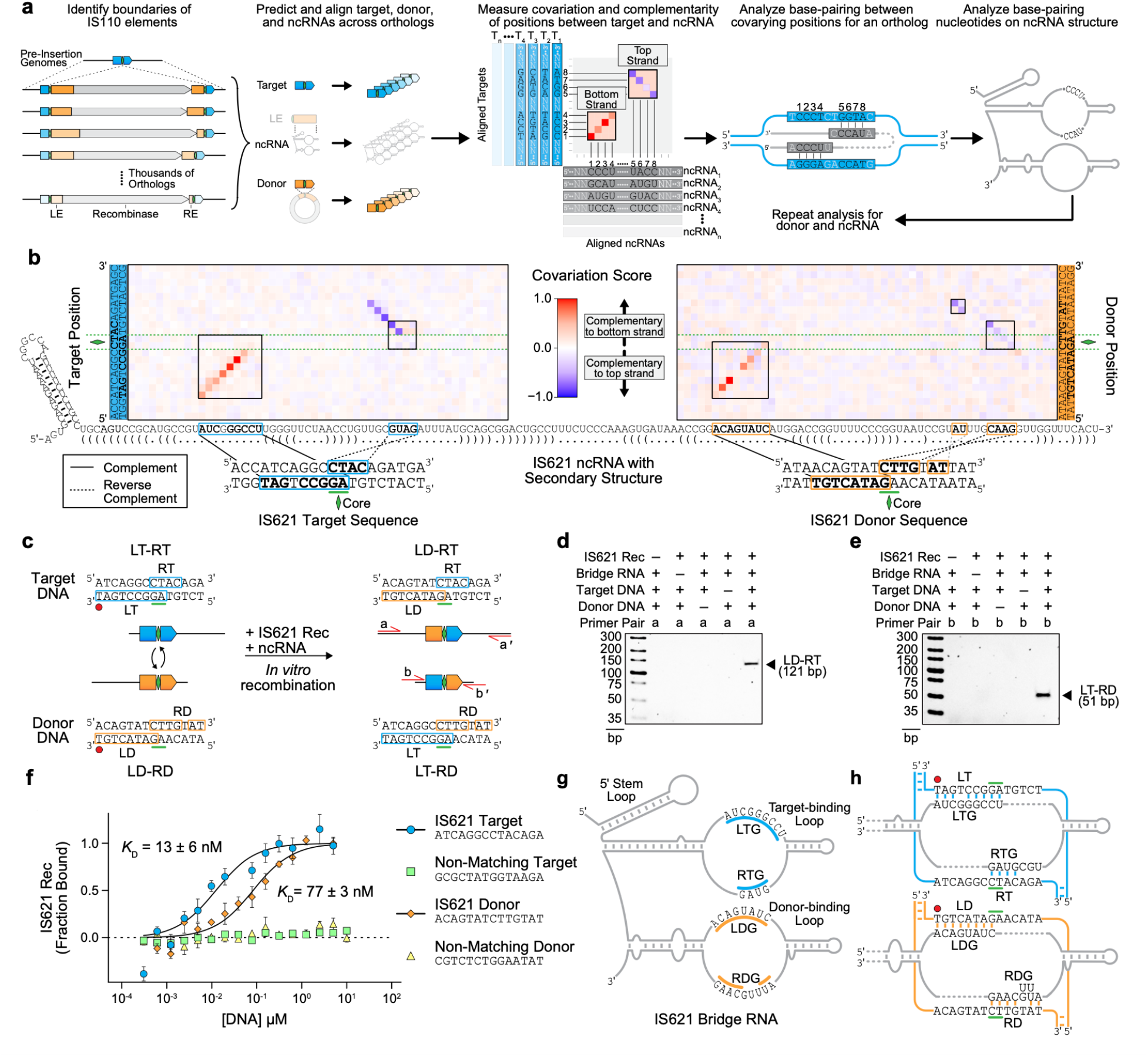
Computational identification of IS621 bridge RNA binding loops with sequence-specific recognition of target and donor DNA. (**a**) Schematic of our computational approach to assess base-pairing potential between the IS110 ncRNA and its cognate genomic target site or donor sequence. Boundaries of thousands of IS110s were used to predict 50-bp target and donor sequences flanking the dinucleotide core (left). A quantification of covariation and base-pairing potential between target-ncRNA or donor-ncRNA pairs yielded a matrix in which predicted base-pairing interactions are depicted by diagonal stretches of red signal (indicating ncRNA complementarity to the bottom strand of the DNA) or blue signal (indicated ncRNA complementarity to the top strand of the DNA) (middle). Consensus regions of covariation were then examined for each IS110 ortholog of interest and mapped onto the predicted ncRNA secondary structure (right). (**b**) Nucleotide covariation and base-pairing potential between the ncRNA and the target (left) and donor (right) sequences across 2,201 ncRNA-target pairs and 5,511 ncRNA-donor pairs, respectively. The canonical IS621 ncRNA sequence is shown across the x-axis, along with dot-bracket notation predictions of the secondary structure. Covariation scores calculated from thousands of IS110 orthologs are colored according to strand complementarity, with -1 (blue) representing high covariation and a bias toward top strand base-pairing, 1 (red) representing high covariation and a bias toward bottom strand base-pairing, and 0 indicating no detectable covariation. Regions of notable covariation signal indicating base-pairing for IS621 are boxed. Blue and orange boxes in the IS621 ncRNA sequence highlight stretches of high base-pairing potential to its native *E. coli* target and donor DNA, respectively. Mapping to the corresponding complementary (solid lines) or reverse complementary (dotted lines) sequences within the double-stranded target DNA (bottom left) or donor DNA (bottom right) is indicated. (**c**) Schematic of *in vitro* recombination (IVR) reaction with IS621. Successful recombination centered at the core dinucleotide (green diamond) would result in two expected products by primer pairs *a* and *b*, respectively. LD, left donor; RD, right donor; LT, left target; RT, right target. (**d**) Gel electrophoresis of IVR LD-RT PCR product. Results are representative of three technical replicates. Rec, recombinase. (**e**) Gel electrophoresis of IVR LT-RD PCR product. Results are representative of three technical replicates. Rec, recombinase. (**f**) Binding of target and donor DNA sequences by IS621 RNP containing fluorescently labeled recombinase and ncRNA. Binding of RNP with cognate or scrambled target and donor DNA sequences is measured using microscale thermophoresis (MST). Mean ± SD for three technical replicates shown. (**g**) Schematic of the IS621 bridge RNA. The target-binding loop contains the left target guide (LTG) and right target guide (RTG) (blue), and the donor-binding loop contains the left donor guide (LDG) and right donor guide (RDG) (orange). (**h**) Base-pairing model of the IS621 bridge RNA with cognate target and donor DNA. The target-binding loop binds both strands of the target (blue), and the donor-binding loop binds both strands of the donor (orange).

To experimentally investigate the potential presence of an IS110-encoded ncRNA, we focused on the IS110 family member IS621, which is native to some strains of *E. coli*, and five closely related orthologs (>76% pairwise amino acid identity). We cloned the concatenated RE-LE sequences of the predicted circular forms onto plasmids and performed small RNA sequencing in *E. coli* BL21(DE3), revealing a continuous peak of RNA sequencing coverage spanning ∼177 bp of the LE starting from the predicted endogenous **𝛔**^70^-like promoter (**Fig. 1e**).

Next, we assessed whether the ncRNA binds the recombinase by measuring the affinity of an *in vitro*-transcribed 177-nt ncRNA from IS621 and its purified cognate recombinase using microscale thermophoresis (MST). We found that the IS621 recombinase binds the ncRNA with high affinity (*K*D = 2.1 ± 0.2 nM), with no binding detected at the concentrations tested of a scrambled and length-matched RNA control (**Fig. 1f**). Our data indicate that IS110 elements can encode a ncRNA in their LE that is produced by an endogenous promoter reconstituted within the circular form of the element. This ncRNA specifically binds to the recombinase enzyme, suggesting that it may play a functional role in recombination.

### Computational analysis predicts that the IS110 ncRNA mediates target- and self-recognition through base-pairing

Evaluating the ncRNA consensus secondary structure across 103 diverse IS110 orthologs revealed a 5′ stem-loop followed by two additional stem-loops with prominent internal loops of similar size (**Fig. 1g**, **Supplementary Fig. 3a-3b**). We observed that the first internal loop has relatively low sequence conservation across orthologs, while the second is much more conserved (**Supplementary Fig. 3c**).

**Figure 3:**
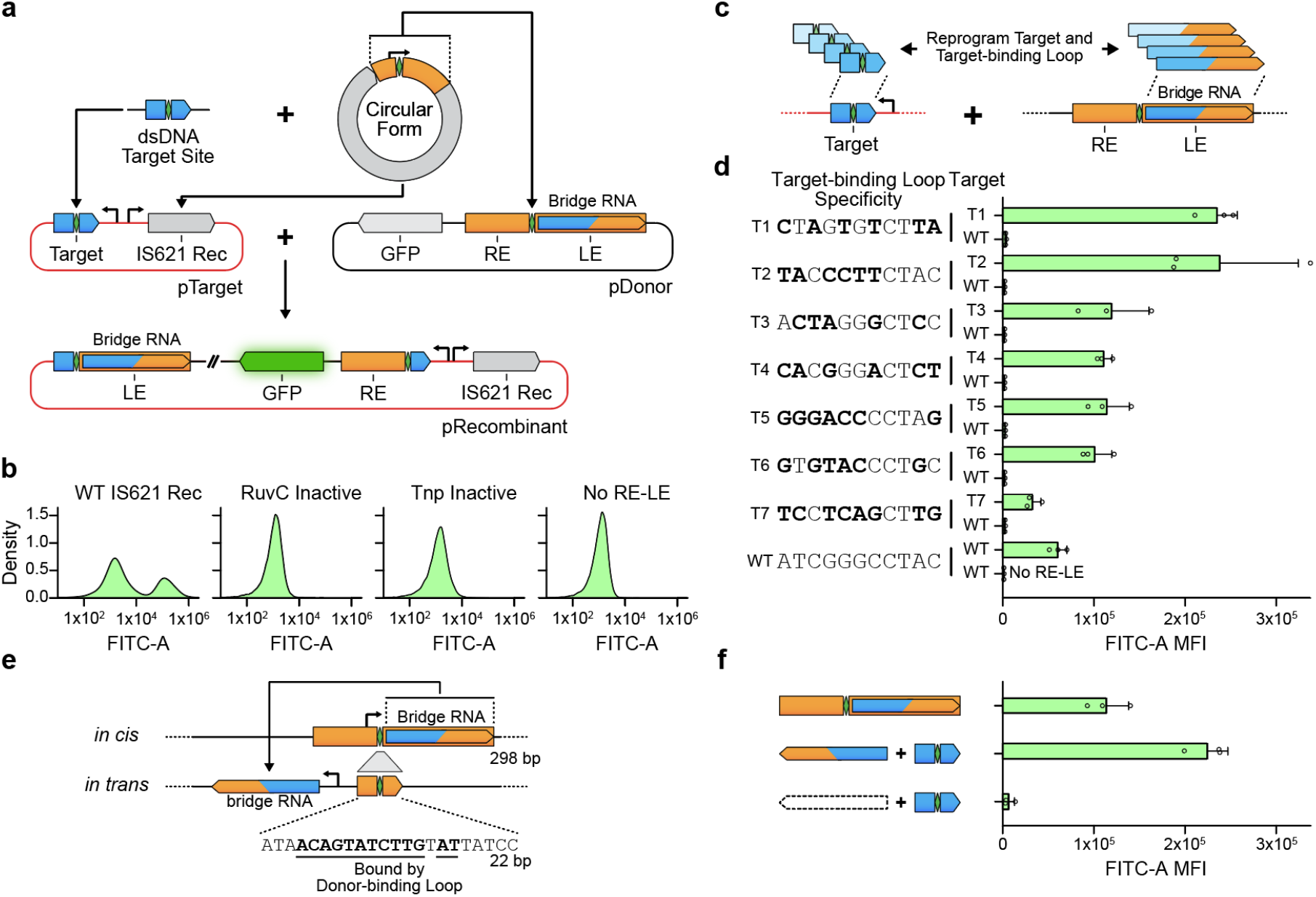
The IS621 target site is reprogrammable and is specified by the bridge RNA. (**a**) Schematic representation of the bacterial recombination assay with bridge RNA *in cis*. Successful recombination centered at the core dinucleotide places GFP downstream of the synthetic promoter, resulting in fluorescence. Rec, recombinase. (**b**) DNA recombination with catalytic variants of IS621 recombinase. From left to right: WT IS621; a variant of the IS621 recombinase containing the D11A/E60A/D102A/D105A mutation in the RuvC active site; a variant of the IS621 recombinase containing the S241A mutation in the Tnp domain active site; and a variant of the pDonor lacking the RE-LE. Plots are representative of 3 replicates. (**c**) Schematic of reprogrammed target and bridge RNA target-binding loop sequences. (**d**) Specific recombination using reprogrammed bridge RNA target-binding loop and target sequences. Target sequences are listed on the left, and the bridge RNA is reprogrammed to base-pair with the indicated sequence. Bold bases highlight differences relative to WT donor sequence. Mean ± SD for three biological replicates shown. (**e**) Schematic of bridge RNA expression *in trans*. Top: Expression of the bridge RNA *in cis* from the natural 𝛔^70^ promoter sequence across the RE-LE junction, as shown in (**a**); Bottom: Expression of the bridge RNA *in trans* from a synthetic promoter; transcriptional termination is achieved via a HDV ribozyme sequence. To eliminate expression of the bridge RNA *in cis* from the donor, the RE-LE junction in the donor is truncated to a 22 bp region lacking the -35 box of the 𝛔^70^ promoter and containing the LD and RD nucleotides required for base-pairing with the bridge RNA donor-binding loop (bold and underlined). (**f**) Comparison of recombination efficiency with bridge RNA expressed *in cis* and *in trans*. Mean ± SD for three biological replicates shown.

Given that the ncRNA can bind the recombinase, we wondered whether it facilitates the recognition of the target site or the donor DNA molecule (i.e. the IS110 element itself) by the recombinase. To assess this, we first systematically defined the element boundaries of thousands of IS110 elements to reconstruct their insertion sites (**Fig. 2a**). Using these boundary predictions, we then reconstructed the target site and circular form for each element. An iterative search using a structural covariance model (CM) that we developed of the IS621 ncRNA enabled the prediction of thousands of ncRNA orthologs encoded within LEs (**Methods**) (Nawrocki & Eddy, 2013).

We then created a paired alignment of these ncRNAs with their respective target and donor sequences: regions which we defined as the 50 bp centered around the target and donor “CT” core, respectively. To assess the possibility of base-pairing between the predicted ncRNAs and their target and donor sequences, we performed a covariation analysis across 2,201 donor-ncRNA pairs and 5,511 target-ncRNA pairs that were detected by homology with the IS621 element (Seemayer et al., 2014) (**Supplementary Information**). Nucleotide sequence covariation would indicate evolutionary pressure to conserve base-pairing interactions between ncRNA positions and target or donor positions. We also incorporated a base-pairing concordance analysis to identify stretches of the ncRNA that might bind with either the top or bottom strand of the target or donor DNA.

This combined analysis indicated a clear potential base-pairing signal between the two internal loops of the ncRNA and the target and donor DNA sequences, respectively (**Fig. 2b**, **Supplementary Fig. 4a-4b**). Projecting this covariation pattern onto the canonical IS621 sequence and ncRNA secondary structure, we inferred that the first internal loop may base-pair with the target DNA, while the second internal loop may base-pair with the donor DNA. The 5′ side of each loop appears to base-pair with the bottom strand of the target/donor with a stretch of 8-9 nucleotides, while the 3′ side of each loop appears to base-pair with the top strand of the target/donor using 4-6 nucleotides (**Fig. 2b**). The strong covariation and base-pairing signal suggested that ncRNA base-pairing with target and donor DNA may be a conserved mechanism across diverse IS110 orthologs.

**Figure 4:**
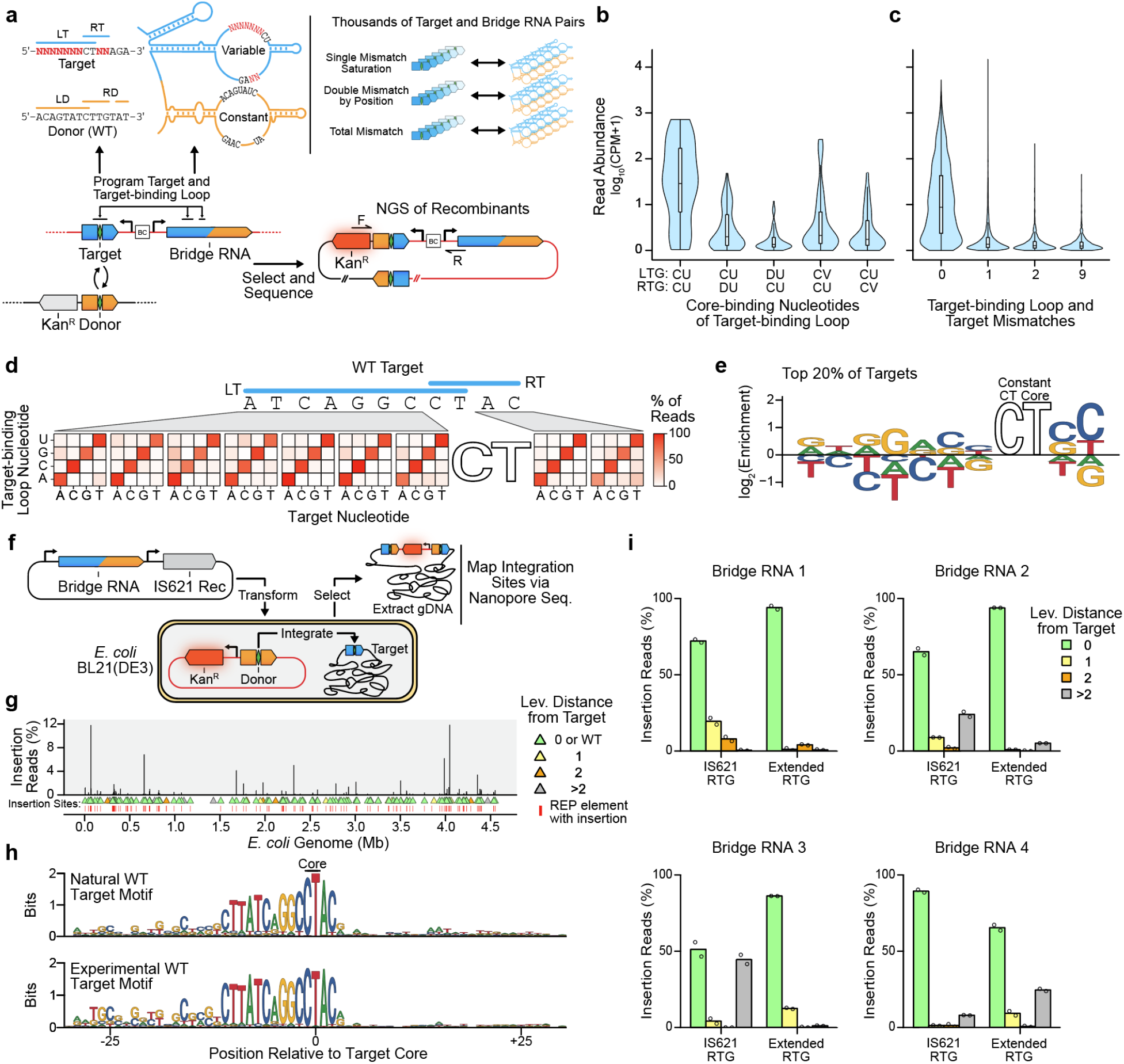
High-throughput characterization of IS621 target specificity demonstrates flexible programmability. (**a**) Schematic representation of a target specificity screen. Successful recombination enables survival of *E. coli* through expression of kanamycin resistance cassette (Kan^R^). The target sequence and bridge RNA are separated by a DNA barcode of 12 nt (BC). A T7-inducible IS621 recombinase is expressed from the plasmid bearing the target and bridge RNA. The WT donor and WT donor-binding loop are used. (**b**) Mismatch tolerance between the target core and target-binding loop. Core-binding nucleotides of the target-binding loop are summarized by IUPAC codes, including D (not C) and V( not U). CPM, counts per million. (**c**) Mismatch tolerance between non-core sequences of the target and target-binding loop. CPM, counts per million. (**d**) Mismatch tolerance between target and target-binding loop by position. The percentage of total detected recombinants encoding each mismatch is shown. The core of the target and core-binding sequences of the target-binding loop are constant across displayed recombinants. (**e**) Nucleotide enrichment of efficient perfectly matched pairs of targets and target-binding loops. The sequence logo depicts the nucleotide enrichment at each position among the top 20% most efficient targets tested in the target specificity screen. (**f**) Schematic of genome insertion assay in *E. coli*. An *E. coli* cell line harbors a replication-incompetent plasmid encoding the WT donor (see Fig. 3e). Integrants are enriched by selecting for expression of kanamycin resistance from the genome after transformation with a plasmid encoding the recombinase and bridge RNA. Integration sites are determined via nanopore sequencing of isolated genomic DNA. (**g**) Genome-wide mapping of insertions mediated by the WT IS621 bridge RNA. The percentage of total reads mapped to each insertion site is depicted. Triangles indicate the number of differences from the intended target site as measured by Levenshtein distance. Insertions into REP elements are highlighted with a red line. (**h**) Insertion sequence preference of IS621. Sequence logos depict the target site motifs among natural (top, **Methods**) and experimentally observed (bottom, see Fig. 4g) IS621 target sites. (**i**) Genomic specificity profile of four reprogrammed bridge RNAs. Using the same approach in (**f**), 11 bp target sequences appearing once in the *E. coli* genome were targeted. The RTG of the bridge RNA was programmed to target the RT of the 11 bp sequence (IS621 RTG, length 4 nt) or the same RT and the 3 adjacent bases (Extended RTG, length 7 nt). Color indicates the number of differences from the intended sites as measured by Levenshtein distance. Data represent sums of all insertion sites with 0, 1, 2 or >2 differences from the intended target.

### The IS621 ncRNA and recombinase catalyze recombination by bridging target and donor DNA

Prior attempts to study or reconstitute IS110-mediated recombination have only been successful in IS110 host organisms, with no reports of successful *in vitro* recombination (Higgins et al., 2007, 2009; Perkins-Balding et al., 1999). We reasoned that the ncRNA could be the missing component required for recombination. To test this, we combined the *in vitro*-transcribed ncRNA with the purified IS621 recombinase and DNA oligonucleotides containing target and donor DNA sequences to assess *in vitro* recombination by PCR and sequencing. Strikingly, we found that the ncRNA is necessary for *in vitro* recombination, and that the four components (ncRNA, recombinase, target DNA, and donor DNA) are sufficient to produce a recombination product containing the expected junction at the core dinucleotide (**Fig. 2c-2e**). MST experiments further revealed that the recombinase-ncRNA ribonucleoprotein (RNP) complex binds wild-type (WT) target and donor DNA oligos (target *KD* = 13 ± 6 nM; donor *KD* = 77 ± 3 nM), but not target and donor molecules containing scrambled sequences at the regions otherwise complementary to the ncRNA (**Fig. 2f**). Taken together, these findings indicate that the ncRNA bound by the IS621 recombinase enables sequence-specific binding to both target and donor DNA molecules, and thereby facilitates recombination.

We named this ncRNA a ‘bridge RNA’, based on its colocalization role in bridging the target and donor DNA molecules for recombination. We refer to the two internal loops of the bridge RNA as the target-binding loop and donor-binding loop (**Fig. 2g**). The target-binding loop comprises two key regions that base-pair with the top and bottom strands of the target DNA, respectively: the left target guide (LTG) base-pairs with the left side of the bottom strand of the target DNA (left target; LT), while the right target guide (RTG) base-pairs with the right top strand of the target DNA (right target; RT). The donor-binding loop has an analogous architecture, with a left donor guide (LDG) base-pairing with the bottom strand of the left donor (LD) and a right donor guide (RDG) base-pairing with the top strand of the right donor (RD) (**Fig. 2h**). Importantly, we noted that the core dinucleotide is included in every one of the base-pairing interactions (LTG-LT, RTG-RT, LDG-LD, and RDG-RD), resulting in an overlap between the right top and left bottom strand pairings and suggesting a key role for bridge RNA-core interactions for recombination.

To lend further support to our model that the bridge RNA target-binding loop guides genomic target site selection, we analyzed natural IS110 insertion loci across diverse IS110 orthologs. Binning natural IS110s by sequence similarity of their LTG and RTG, we created a consensus genomic target site motif for each LTG/RTG. The target motif was highly concordant with the target-binding loop sequences of the bridge RNA (LTG and RTG), with 0-2 mismatches in most cases (**Fig. 2b**, **Supplementary Fig. 5a**). Through further inspection of our covariation data, we noticed that the RTG of some IS110 orthologs may be longer than the RTG for IS621 (**Fig. 2b**, **Supplementary Fig. 5b-5c**). We also observed evidence of an unusual base-pairing pattern between the RDG and the RD, where a stretch of 9 bridge RNA bases base-pairs discontiguously with a stretch of 7 donor DNA bases (**Fig. 2b**, **Fig. 2h**).

**Figure 5:**
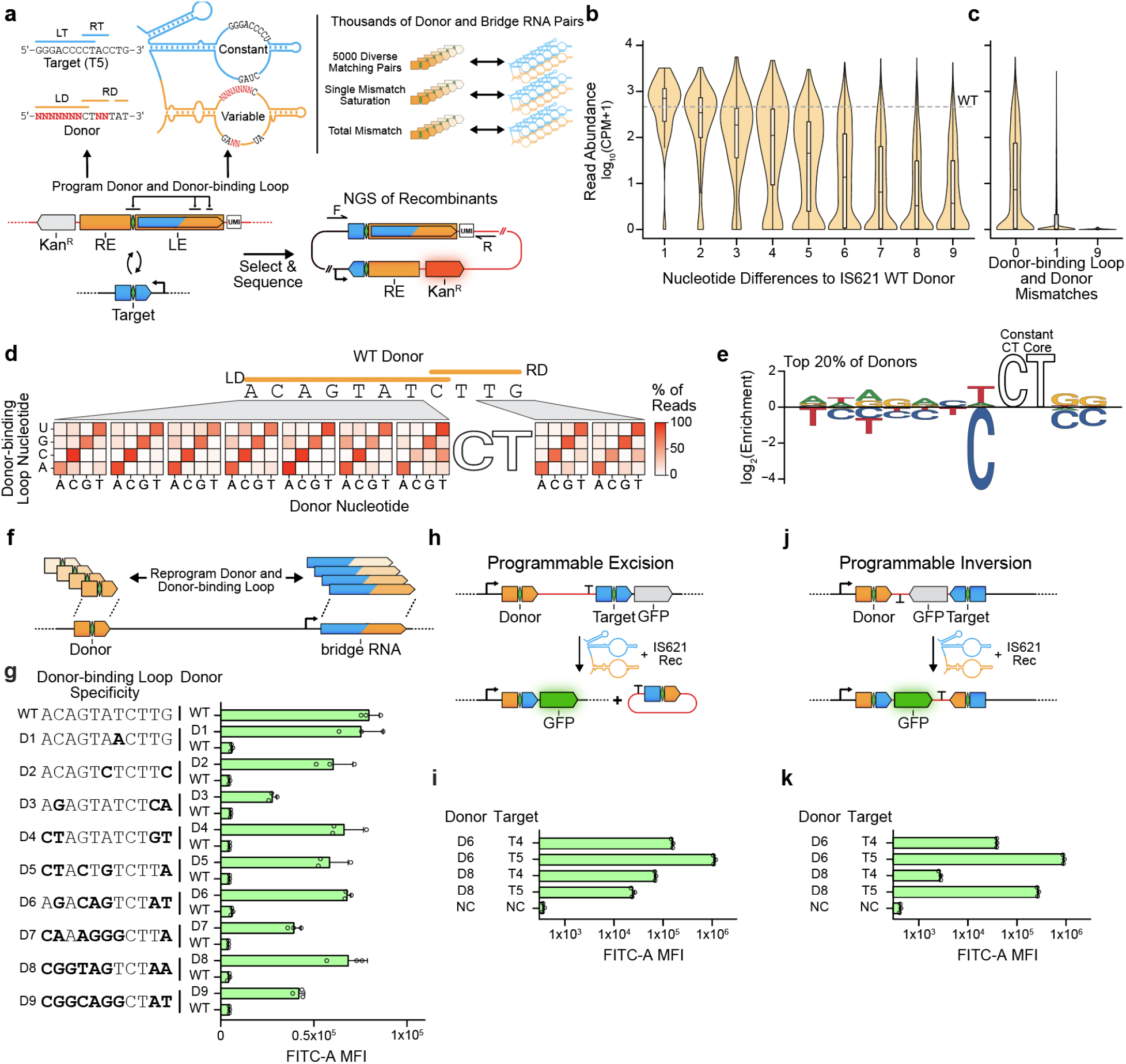
Bridge RNA donor recoding enables fully programmable insertion, inversion, and excision. (**a**) Schematic representation of the high throughput donor screen. The recombination assay design uses the same approach as in Fig. 3a and 3c, except that the expression of a kanamycin resistance gene is induced and surviving colonies are assayed. A unique molecular identifier (UMI) identifies each pair of donor and donor-binding loops. The bridge RNA target-binding loop is matched to target T5. Two replicates were averaged to produce read abundance per oligo, quantified as CPM, counts per million. NGS, next generation sequencing. (**b**) Reprogrammability of donor sequences by number of nt differences from the WT donor. Wild-type donor abundance is indicated by the gray dashed line. CPM, counts per million. (**c**) Mismatch tolerance between non-core sequences of the donor-binding loop and donor. CPM, counts per million. (**d**) Mismatch tolerance between bridge RNA donor-binding loop and donor by position. The percentage of total detected recombinants encoding each single mismatch is recorded. The core of the donor and core-binding sequences of the donor-binding loop are constant across displayed recombinants. (**e**) Nucleotide enrichment of efficient perfectly matched pairs of donors and donor-binding loops. Sequence logo represents the nucleotide enrichment at each position among the top 20% most efficient targets tested in the donor specificity screen. (**f**) Schematic illustrating paired reprogramming of donor/donor-binding loop. donor-binding loops were reprogrammed and tested against the wild-type donor and their cognate donor. (**g**) Specific recombination using reprogrammed donor and bridge RNA donor-binding loop sequences. Donor sequences are listed at left, and the bridge RNA is reprogrammed to base-pair with the indicated sequence. Bold bases highlight differences relative to WT donor sequence. MFI ± SD for three biological replicates shown. (**h**) Schematic representation of the programmable excision assay. (**i**) Efficient programmable excision of DNA. Pairs of donor and target are denoted. Negative control (NC) expresses the reporter and recombinase but no bridge RNA. MFI ± SD for three biological replicates shown. (**j**) Schematic representation of the programmable inversion recombination assay. (**k**) Efficient programmable inversion of DNA. Pairs of donor and target are denoted. Negative control (NC) expresses the reporter and recombinase but no bridge RNA. MFI ± SD for three biological replicates shown.

### The IS621 bridge RNA can be reprogrammed to direct insertion into diverse target sites

We reasoned that the base-pairing mechanism of target and donor recognition suggested by our computational analysis indicates that the bridge RNA sequence can be altered to reprogram IS621 target and/or donor specificity. To assess this, we set up a two-plasmid recombination reporter system in *E. coli*. pTarget encodes the IS621 recombinase along with a 50-bp target site flanked by a promoter. pDonor contains the concatenated RE-LE donor sequence encoding the bridge RNA as well as a promoter-less *gfp* cargo gene. Recombination of pDonor into the target site on pTarget would place the *gfp* ORF downstream of the promoter, allowing for detection of successful recombination events using flow cytometry (**Fig. 3a**). Using the WT IS621 donor and target sequences in this assay, we detected GFP expression and confirmed the expected recombination product using nanopore sequencing (**Fig. 3b**). Alanine substitution of the conserved catalytic residues of either the RuvC-like domain or the Tnp domain (**Supplementary Fig. 1a-1b**) abolished recombination, as did substituting pDonor with a version lacking the RE-LE (and therefore lacking the bridge RNA) (**Fig. 3b**).

In addition to the WT target and WT bridge RNA sequence used above, we selected seven target sequences (T1-T7) that were orthogonal to the *E. coli* genome and designed reprogrammed bridge RNAs with matching target-binding loops according to the base-pairing rules we derived from our covariation analysis (**Fig. 3c**). Reprogramming the bridge RNA target-binding loops to match targets T1-T7 prevented recombination with the non-matching WT target in all cases, while enabling high rates of recombination (13.8-59.5% of all cells) with its cognate target sequence (**Fig. 3d**, **Supplementary Fig. 6a**). We next asked whether the bridge RNA could be expressed in *trans* rather than within the RE-LE context. First, we truncated the RE-LE (298 bp) to a 22 bp donor around the core dinucleotide to maintain a functional donor sequence but eliminate the -35 box of the natural 𝛔^70^ promoter as well as the majority of the bridge RNA-encoding sequence (**Fig. 1e**). Using this variant of pDonor, we observed no recombination between the truncated donor and the T5 target plasmid. We next added the full-length bridge RNA-encoding sequence elsewhere on pDonor under the control of a synthetic promoter. Expression of the T5-specific bridge RNA in this context not only restored recombination of pDonor with pTarget-T5, but did so more robustly than the same bridge RNA expressed from the endogenous promoter, increasing FITC-A signal by 1.98-fold. (**Fig. 3e-f**). Taken together, these results indicate that the bridge RNA target-binding loop can be reprogrammed in *cis* or in *trans* to direct target site specificity for DNA recombination in *E. coli*.

**Figure 6:**
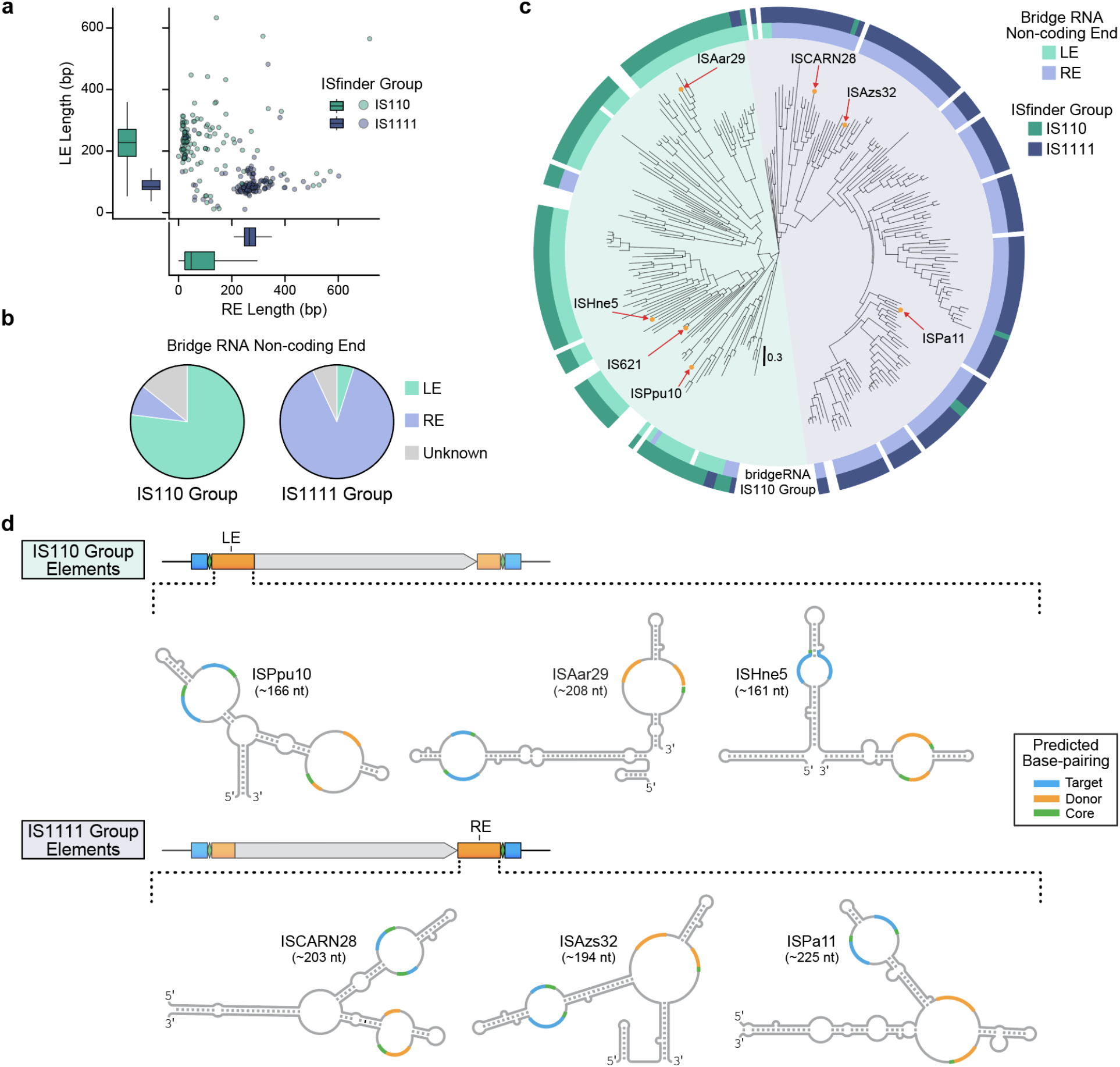
IS110 subfamilies encode distinct and diverse bridge RNA secondary structures in different non-coding end sequences. (**a**) Non-coding end length distribution for IS110 and IS1111 group elements. (**b**) Location of predicted bridge RNA for IS110 and IS1111 group elements. (**c**) Phylogenetic tree of the 274 IS110 recombinases cataloged by ISFinder. (**d**) Bridge RNA consensus structures from six diverse IS110 elements. Secondary structures are shown with internal loops colored according to the sequence that they complement – target (blue), donor (orange), or core (green). Three members of each IS110 group are shown.

### Systematic screen of the bridge RNA target-binding loop demonstrates specific and broadly reprogrammable recombination

To comprehensively assess bridge RNA mismatch tolerance and reprogramming rules, we designed an *E. coli* selection screen that links thousands of barcoded pairs of DNA targets and bridge RNAs on a single plasmid. Successful recombination with a WT donor plasmid induces a kanamycin resistance cassette (Kan^R^) for survival (**Fig. 4a**). Using this approach, we first confirmed that base-pairing between the bridge RNA and both strands of the CT target core sequence was strongly preferred, in line with the high conservation of the CT core sequence in both the target and donor (**Fig. 4b**, **Supplementary Fig. 5d-5e**, **Supplementary Fig. 6b**).

Next, we varied the nine non-core positions of the target and the corresponding positions of the LTG and RTG, intending to assess single and double mismatch tolerance at each position. We observed a strong preference for perfect matches across all nine positions of the target-binding loop and target, and a high degree of flexibility to full reprogramming of all positions (**Fig. 4b-4d**, **Supplementary Fig. 6b** and **6c**). As expected, double mismatches were even less well tolerated than single mismatches, with bias for certain mismatch position combinations (**Fig. 4c**, **Supplementary Fig. 6c**). Overall, although we observed some position-specific nucleotide preferences, our screen demonstrated that the target-binding loop is broadly programmable at each position with low mismatch tolerance (**Fig. 4e**).

### Bridge RNA-mediated integration of large cargos into the *E. coli* genome

To evaluate genomic site-selection and specificity of WT IS621, we measured integration of a replication-incompetent plasmid (4.85 kb) bearing the 22 bp WT donor sequence into the *E. coli* genome using the WT IS621 bridge RNA and recombinase (**Fig. 4f**). After selecting for integration using kanamycin, we mapped insertions genome-wide in *E. coli* at >1000x colony coverage and observed 173 unique insertion sites, with 144 of these insertions occurring within the predicted REP elements known to be naturally targeted by WT IS621 (**Fig. 4g**) (Choi et al., 2003). Of all insertion sites, 74.5% (129 sites) matched the WT target sequence (ATC**A**GGCCTAC), while two more sites exactly matched the specificity encoded by the target binding loop (ATC**G**GGCCTAC); these two target sequences accounted for 96.21% of detected insertions (**Supplementary Fig. 7a-c**). Accordingly, our experimental results closely mirrored the specificity profile of IS621 elements found in nature, including tolerance for a mismatch at position 4 of the target site (**Fig. 4h**). Further analysis of the IS621 recombinase:bridge RNA complex bound to target DNA indicated that this perceived mismatch results in a non-canonical rG:dT base pair, potentially explaining the high frequency of insertions into the WT target sites.

Based on our target specificity screen, we expected that the two perfectly matching sequences in the genome would be the most frequently targeted insertion sites. However, we found that 13 sites were more frequently selected. Further scrutiny of these insertion sites revealed that four of the ten most frequently targeted sites were flanked on the 3′ end of the RT sequence by 5′-GCA-3′ – complementary to the 5′-UGC-3′ that occurs immediately 5′ of the RTG in the WT bridge RNA (**Fig. 2b**, **Supplementary Fig. 7a-7d**). This suggested to us the potential of an extended base-pairing interaction beyond the predicted RTG:RT for IS621 (7 bp instead of 4 bp), which was further supported by the observation in our covariation analysis that some IS110 orthologs naturally encode longer RTGs (**Fig. 2b**, **Supplementary Fig. 5b-5c**).

To investigate genome-wide specificity of insertion into reprogrammed sites, we designed bridge RNAs to target unique sites found only once in the *E. coli* genome. We tested four distinct genomic sites with two bridge RNAs for each: one harboring a short 4 bp RTG (IS621 RTG) and one with a long 7 bp RTG (Extended RTG) to directly assess the impact of RTG:RT base-pairing length on specificity and/or efficiency. In every case, we found that the expected genomic target site was the most frequently targeted, representing between 51.6% and 94.0% of all detected insertions (**Fig. 4i**). Off-target insertions were also observed, with individual off-target sites representing between 0.11% and 31.16% of insertions across all bridge RNAs, with the more frequently detected off-targets typically carrying 1-2 mismatches with the expected target (**Supplementary Fig. 7e**).

The extended RTG improved the frequency of insertion into the on-target site from an average of 69.4% (range 51.2%-89.4%) to an average of 84.9% (range 65.4%-94.0%). It also resulted in significantly fewer insertions into off-target sites for bridge RNA 2 and bridge RNA 3, eliminating 18/45 and 14/25 off-target sites, respectively (**Fig. 4i**). We found further support for the longer RTG-RT interaction improving efficiency from our analysis of off-targets for bridge RNA 1: two of the higher-frequency off-targets for the 4 bp IS621 RTG exhibited a natural base-pairing extension (flanked by 5′-GCA-3′), similar to what we had observed for enriched insertion sites for the WT IS621 bridge RNA (**Supplementary Fig. 7d**). Intriguingly, some off-target sites seemed to indicate tolerance for insertions in the target sequence, while some low-frequency insertions appeared to more closely resemble the 11 bp WT donor sequence rather than the programmed target (**Supplementary Fig. 7e-f**).

Of the 117 genomic off-target insertion sites detected across the 8 experiments, 102 (87.2%) had the expected CT dinucleotide core, 56 (47.9%) closely resembled the target or donor sequence (Levenshtein distance < 3), and the remaining sites were enriched for long *k*-mer matches to the target or donor sequence (**Supplementary Fig. 7g**), suggesting most or all detected off-target insertions were bridge RNA-dependent. In addition to off-target insertions, genomic deletions and inversions between experimentally observed insertion sites were observed with rare frequency (allele frequency <0.05) (**Supplementary Note 1**). Altogether, these experiments demonstrate the robust capability of IS621 to specifically integrate large, multi-kilobase cargos into the genome in a reprogrammable fashion and offer further insights into the recombination mechanism.

### Bridge RNA donor-binding loops can be broadly reprogrammed for specific recombination

In naturally occurring IS621 elements, the donor sequence is much more highly conserved than the genomic target sequence, suggesting that the donor-binding loop may be less readily reprogrammed than the target-binding loop (**Supplementary Fig. 5d-5e**). To assess this, we designed a donor specificity screen in which we varied the 7-bp LD and 2-bp RD flanking the core dinucleotide, all within the context of a full-length RE-LE expressing the bridge RNA in *cis*. Successful recombinants with the T5 target plasmid would induce Kan^R^ expression and enable *E. coli* survival (**Fig. 5a**). Analysis of thousands of matching donor and donor-binding loop pairs revealed that despite conservation of the WT donor sequence, it can be reprogrammed with any number of differences from the WT donor (**Fig. 5b**). Similar to the interaction between the target and target-binding loop of the bridge RNA, LD-LDG mismatches and RD-RDG mismatches were generally poorly tolerated (**Fig. 5c**). Position 7 of the LD (directly flanking the core dinucleotide) was an exception to these results, exhibiting a strong bias against cytosine and therefore appearing more mismatch tolerant than other positions (**Fig. 5d-5e**).

In these experiments, the core dinucleotide (CT) was held constant, which could limit the sequence space of potential target and donor sites. To address this, we modified the conserved cores of target T5 and the WT donor, along with their associated bridge RNA positions in both loops, from CT to AT, GT, or TT (**Supplementary Fig. 8a-b**). We found that, while non-CT cores were generally less efficient, efficiency was improved by extending the length of RTG-RT base-pairing from 4 to 7 bp, informed by our earlier results on RTG extension (**Fig. 4i**, **Supplementary Fig. 8c-d**)

Next, we explored the ability of the bridge RNA to combinatorially control the recognition of target and donor sequences simultaneously. Utilizing our *trans* GFP reporter assay where the target-binding loop of the bridge RNA recognizes target T5 (**Fig. 3e**), we reprogrammed the donor sequence and the donor-binding loop of the bridge RNA to one of nine distinct donor sequences (D1-D9) with varying levels of sequence divergence from the WT donor (**Fig. 5f**). Pairing the distinct donor-binding loops with either the reprogrammed donor or the WT donor, we found that matching donor-binding loop and donor sequences that are highly divergent from the WT donor can result in efficient and specific recombination, even when both target and donor are distant from the WT sequences.(**Fig. 5g**). Taken together, we demonstrate that the bridge RNA allows modular reprogramming of both target and donor recognition.

### Programmable rearrangement of DNA sequences in *E. coli*

In addition to their use for DNA insertion, established recombinases such as Cre have been routinely used for excision or inversion of a fixed DNA sequence within a single molecule. Typically, such approaches require pre-installation of the loxP recognition sites in the appropriate arrangement, with two sites oriented in the same direction resulting in excision, and sites oriented in opposite directions resulting in inversion. Given our understanding of the IS621 insertion mechanism, as well as the reported existence of invertase homologs of IS110s (Rozsa et al., 1997; Tobiason et al., 2001), we hypothesized that IS621 recombinases could mediate excision and inversion in a similar way to Cre, but with programmable recognition sites.

We first generated GFP reporter systems for both excision and inversion, in which GFP expression would only be induced after the desired rearrangement occurred (**Fig. 5h-5k**, **Supplementary Fig. 9a-9c**). Testing the same four pairs of donor and target recognition sites in both reporters, we demonstrated that both reactions occur robustly and in a reprogrammable manner. Overall, the ability of IS110 recombinases and their bridge RNAs to insert, excise, and invert DNA in a programmable and site-specific manner enables remarkable control over multiple types of DNA rearrangements with a single unified system.

### Diverse IS110 elements encode programmable bridge RNAs

Our findings for IS621 led us to explore whether the bridge RNA is a general feature of the IS110 family. The IS110 family is divided into two groups: one named IS110 (which includes IS621) and the other called IS1111. IS1111 elements also encode DEDD recombinases, but have been categorized into a separate group based on the presence of sub-terminal inverted repeat sequences (STIRs) that range in length from 7 to 17 bp (Müller et al., 2001; Partridge & Hall, 2003; Siguier et al., 2015). We closely examined our covariation analysis of IS110 termini and identified a short 2-3 bp STIR pattern that flanks the programmable donor sequence, suggesting a possible evolutionary relationship with the longer STIRs of IS1111 elements (**Supplementary Fig. 10a**). Amongst all IS110 and IS1111 elements annotated in the ISfinder database (274 elements after 90% amino acid identity clustering), we found that IS1111 elements had a much longer RE than LE, in contrast to the IS110 subgroup where the LE is significantly longer than the RE (**Fig. 6a**).

Using RNA structural covariance models, we detected a predicted bridge RNA in 85.7% of IS110s and 93.0% of IS1111s (**Fig. 6b**). The vast majority of IS110 group members appeared to encode a bridge RNA within the LE while IS1111 appeared to encode a bridge RNA within the RE, consistent with a previous report that correlated target site preference to sequence conservation in the RE of IS1111 elements (Post & Hall, 2009). Remarkably, the location of the bridge RNA closely predicted the phylogenetic relationship between IS110 and IS1111, strongly suggesting that these two groups emerged from a common ancestor where the bridge RNA translocated between the ends of the element and the length of the STIR was modified (**Fig. 6c**).

Having characterized the target and donor binding patterns and programmability of IS621, we aimed to extend these insights across IS110 and IS1111 orthologs. We predicted bridge RNA structures and manually inspected the loops of 6 elements (each sharing less than 25% recombinase amino acid identity) for evidence of complementarity with the target and donor sequences. Diverse structures emerged from this analysis, with clear evidence of a base-pairing pattern between internal bridge RNA loops and DNA targets and donors (**Fig. 6d**, **Supplementary Fig. 10b**). Interestingly, in many IS1111 orthologs, the predicted bridge RNA has potential donor-binding nucleotides in a multi-loop structure rather than in a simple internal loop for IS621 and other IS110 group members. Altogether, we conclude that the IS110 family encodes diverse predicted bridge RNAs that direct sequence-specific and programmable recombination between target and donor sequences.

## DISCUSSION

Non-coding RNA molecules that bind a specific nucleic acid target are central to both prokaryotic and eukaryotic life. Nucleic acid guides are a widely used mechanism in fundamental biological processes, such as tRNA anticodons that govern ribosomal translation, small interfering RNAs and microRNAs of RNA interference, crRNAs of CRISPR-Cas immunity, and small nucleolar RNAs for gene regulation. To enable the modular control of enzymatic target specificity, their common theme involves a fixed interaction between a guide RNA and an effector protein along with a variable nucleic acid target site specified by base-pairing with the guide.

The bridge RNA we discovered in this work is the first example, to our knowledge, of a bispecific guide molecule that encodes modular regions of specificity for both the target and donor DNA, coordinating these two DNA sequences in close proximity to catalyze efficient recombination. Bridge RNAs encode all of this complex molecular logic in a remarkably compact (∼150-250 nt) sequence along with their single effector recombinase (∼300-460 aa) partner. IS110 targeting is achieved using internal binding loops reminiscent of tRNA hairpin loops or snoRNA internal loops, distinct from the terminal binding sequences of CRISPR-Cas or Argonaute guide RNAs. Each RNA loop encodes segments that base-pair with staggered regions of the top and bottom strand of each cognate DNA binding partner, in contrast to the single-strand base-pairing mechanisms of known RNA-guided systems. Finally, the RNA-guided self-recognition of the IS110 element in donor form illustrates a previously unobserved mechanism of DNA mobility.

Mobile genetic elements are professional DNA-manipulating machines that have been shaped throughout evolution to insert, excise, invert, duplicate, and otherwise rearrange DNA molecules. Bridge RNAs enable IS110 recombinases to leverage the inherent logic of RNA-DNA base-pairing, directly bypassing the complex target site recognition codes of other known transposases and recombinases, which depend on extensive protein-DNA or short ssDNA-DNA interactions that offer much less opportunity for straightforward programmability (Barabas et al., 2008; Buchholz & Stewart, 2001; Lansing et al., 2020; Morero et al., 2018). The IS110 family is evolutionarily diverse and widespread across bacterial and archaeal phyla, providing a rich landscape for further functional and structural insights. In our initial survey of diverse IS110 orthologs, we uncovered a variety of bridge RNA structures and lengths, suggesting significant mechanistic diversity both between and within each of the two major IS110 and IS1111 subfamilies.

Guide RNAs are underpinning a technological revolution in programmable biology (Fire et al., 1998; Jinek et al., 2012; Cong et al., 2013; Hsu et al., 2014; Konermann et al., 2018; Anzalone et al., 2019). However, the direct enzymatic activity of standalone, naturally occurring reprogrammable RNA-guided proteins has been notably limited to the endonuclease function (Jinek et al., 2012; Meister et al., 2004). Successive generations of programmable nucleases and nickases have advanced the prevailing genome editing method from the original homology-based capture of a DNA donor (Capecchi, 1989) to the targeted stimulation of donor insertion, all of which require a complex interplay with endogenous DNA repair processes (Urnov et al., 2010; Cong et al., 2013; Komor et al., 2016; Anzalone et al., 2019). Functional diversification of these systems beyond nucleic acid binding or cleavage has generally required the recruitment or fusion of additional effector proteins, resulting in increasingly large and intricate engineered genome editing fusions (Anzalone et al., 2020; Tou & Kleinstiver, 2022).

The IS110 bridge system, in contrast, employs a single and compact RNA-guided recombinase protein that is necessary and sufficient for direct DNA recombination (**Fig. 2d-e**). Modular reprogramming of donor and target recognition by the bispecific bridge RNA uniquely enables the three fundamental DNA rearrangements of insertion, excision, and inversion for manipulating large-scale DNA sequences and overall genome organization. With further exploration and development, we expect that the bridge mechanism will spur a third generation of programmable RNA-guided tools beyond RNAi and CRISPR-based mechanisms to enable a new frontier of genome design.

## SUPPLEMENTARY FIGURES

**Supplementary Figure 1:**
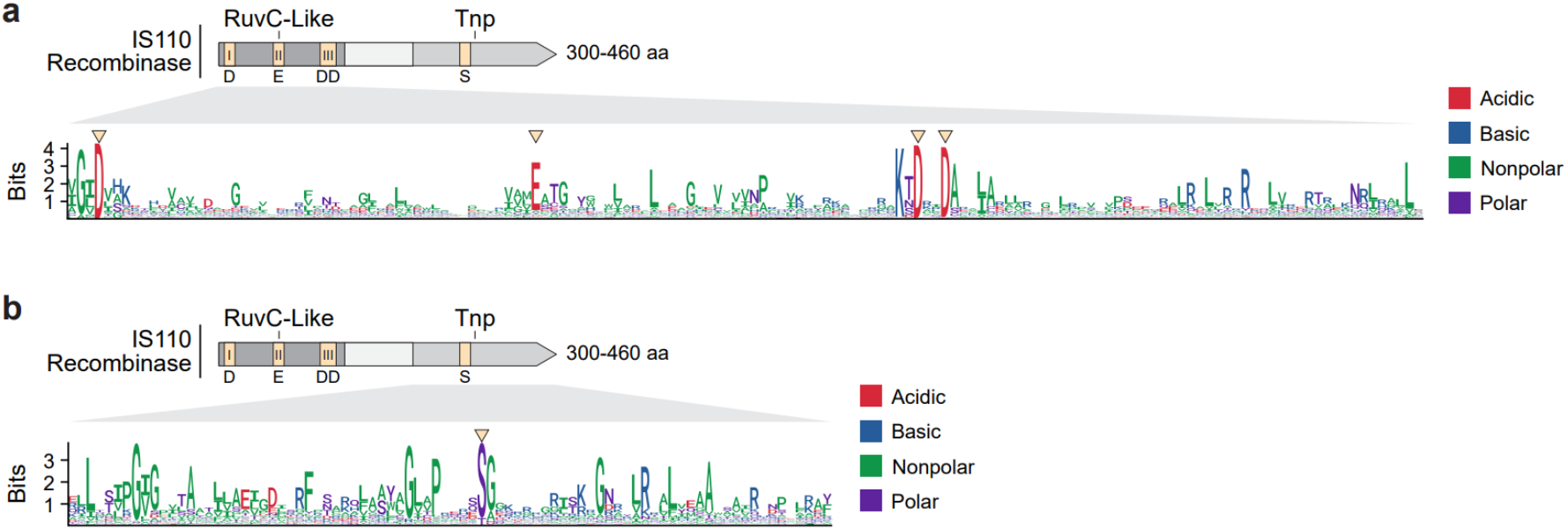
Conserved residues in the RuvC-like and Tnp domains. (**a**) Sequence logo of 213,171 aligned RuvC-like domains identified in IS110 protein sequences. The RuvC-like and Tnp domains shown here were identified using hmmsearch and Pfam models DEDD_Tnp_IS110 (PF01548.19) and Transposase_20 (PF02371.18), respectively. RuvC-like domains were aligned using hmmalign, and these alignments were visualized to identify conserved residues. The conserved residues of the characteristic DEDD motif are highlighted with an arrowhead. The y-axis indicates entropy at each position as measured in bits, with log220 ≈ 4.32 bits being maximally conserved. (**b**) Sequence logo of 208,634 aligned Tnp domains identified in IS110 protein sequences. The Tnp domains were identified, extracted, and analyzed using the same procedure as for the RuvC-like domain. A highly conserved serine residue is highlighted with an arrowhead. The y-axis indicates entropy at each position as measured in bits, with log220 ≈ 4.32 bits being maximally conserved.

**Supplementary Figure 2:**
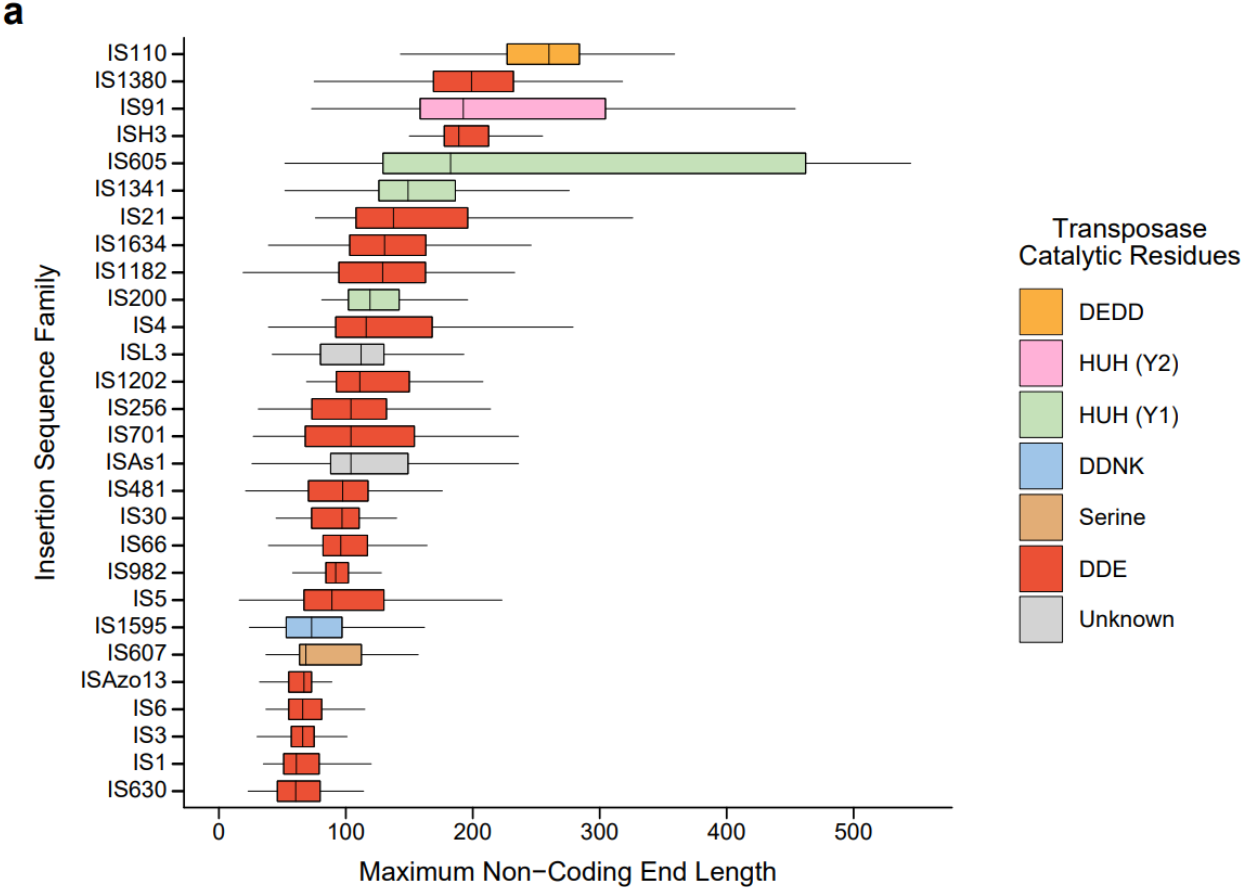
Maximum non-coding end length distribution of 28 insertion sequence families. (**a**) Distribution of non-coding end lengths across different IS families. Boxplot depicting the distribution of non-coding end lengths across IS families, calculated using maximum RE-LE length of 90% identity clusters. Showing 1st and 3rd quartiles (hinges), median, and whiskers extending to 1.5x interquartile range (IQR).

**Supplementary Figure 3:**
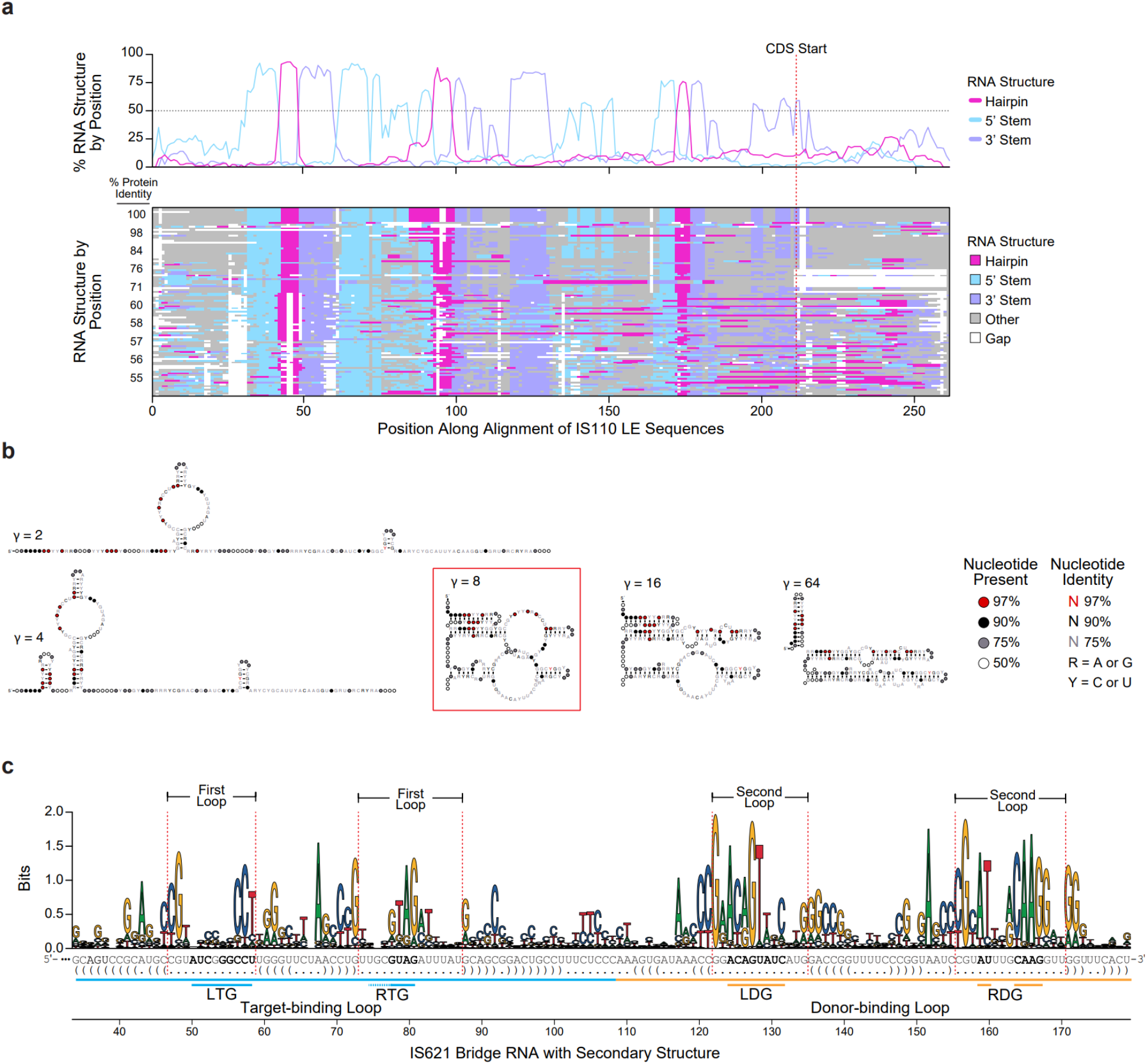
Secondary structure alignment of IS621 non-coding ends and consensus secondary structure prediction. (**a**) Secondary RNA structure alignment of the LE of 103 orthologs of IS621. Secondary RNA structures of the LE of 103 orthologs are predicted and aligned by cluster identity. The percentage of each position corresponding to a 5′ stem, hairpin, or 3′ stem are plotted with a dotted line indicating structures that are conserved in over 50% of sequences. For LE sequences shown along the y-axis, the similarity of their cognate proteins relative to the IS621 recombinase is indicated. This type of visualization was often used throughout the study to determine the presence or absence of a structured ncRNA sequence in the flanks of IS110 recombinase ORFs. (**b**) RNA structures predicted from the LE sequence alignment in (A). RNA structures were predicted using ConsAliFold, which uses a parameter γ to control the prediction balance between positive values (or sequence alignment column base-pairings) and negative values (or unpaired sequence alignment columns). Higher values of γ result in more predicted base-pairing. Showing structures resulting from γ = 2, γ = 4, γ = 8, γ = 16, and γ = 64. The value γ = 8 was used for the initial IS621 ncRNA model in this study. (**c**) Nucleotide conservation across the predicted ncRNA. 2,715 ncRNA orthologs sequences were identified using an iterative search with the original IS621 model, and then aligned with cmalign. The x-axis indicates conservation of nucleotides as measured in bits, quantifying entropy. Highlighting the regions within the prominent internal loops with dotted red lines, with dot-bracket RNA secondary structure notation along the x-axis. The first loop has low sequence conservation (average information content = 0.48 ± 0.09), while the second one is much more conserved (average information content = 0.93 ± 0.11). Sequence features of the bridge RNA are highlighted for clarity.

**Supplementary Figure 4:**
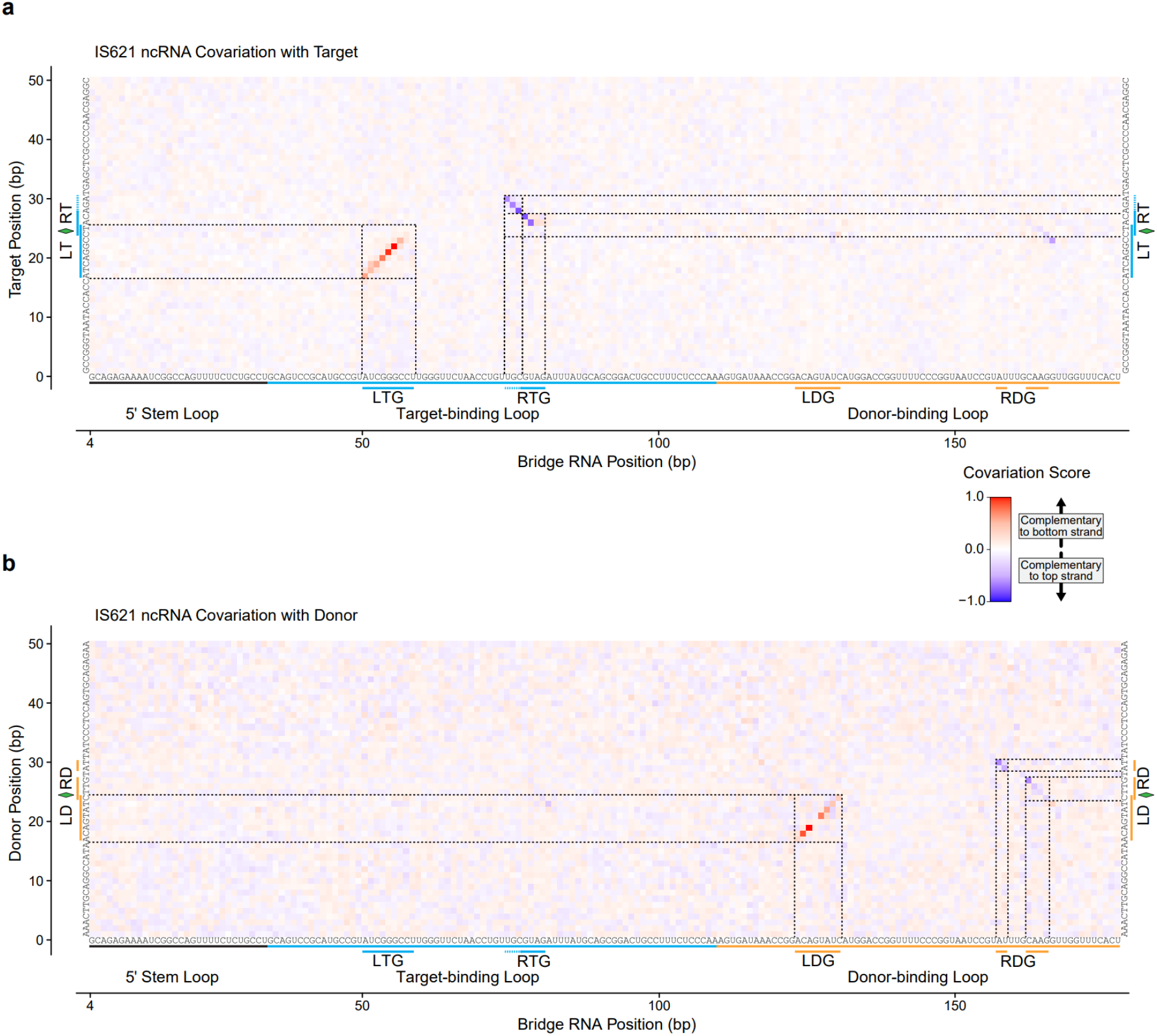
Full ncRNA, target, and donor sequence covariation analysis. (**a**) Expanded schematic of the target-ncRNA covariation data, zoomed out to include the full target and ncRNA sequences used in the analysis. The corresponding IS621 ncRNA sequence and target used in this study is projected along the x-axis. Covariation scores are colored according to the sign (+1 or -1) of the column-permuted base-pairing score to better visualize base-pairing patterns. Key features of the bridge RNA are highlighted for clarity. (**b**) Expanded schematic of the donor-ncRNA covariation data, zoomed out to include the full donor and ncRNA sequences used in the analysis. Features of the bridge RNA are highlighted for clarity.

**Supplementary Figure 5:**
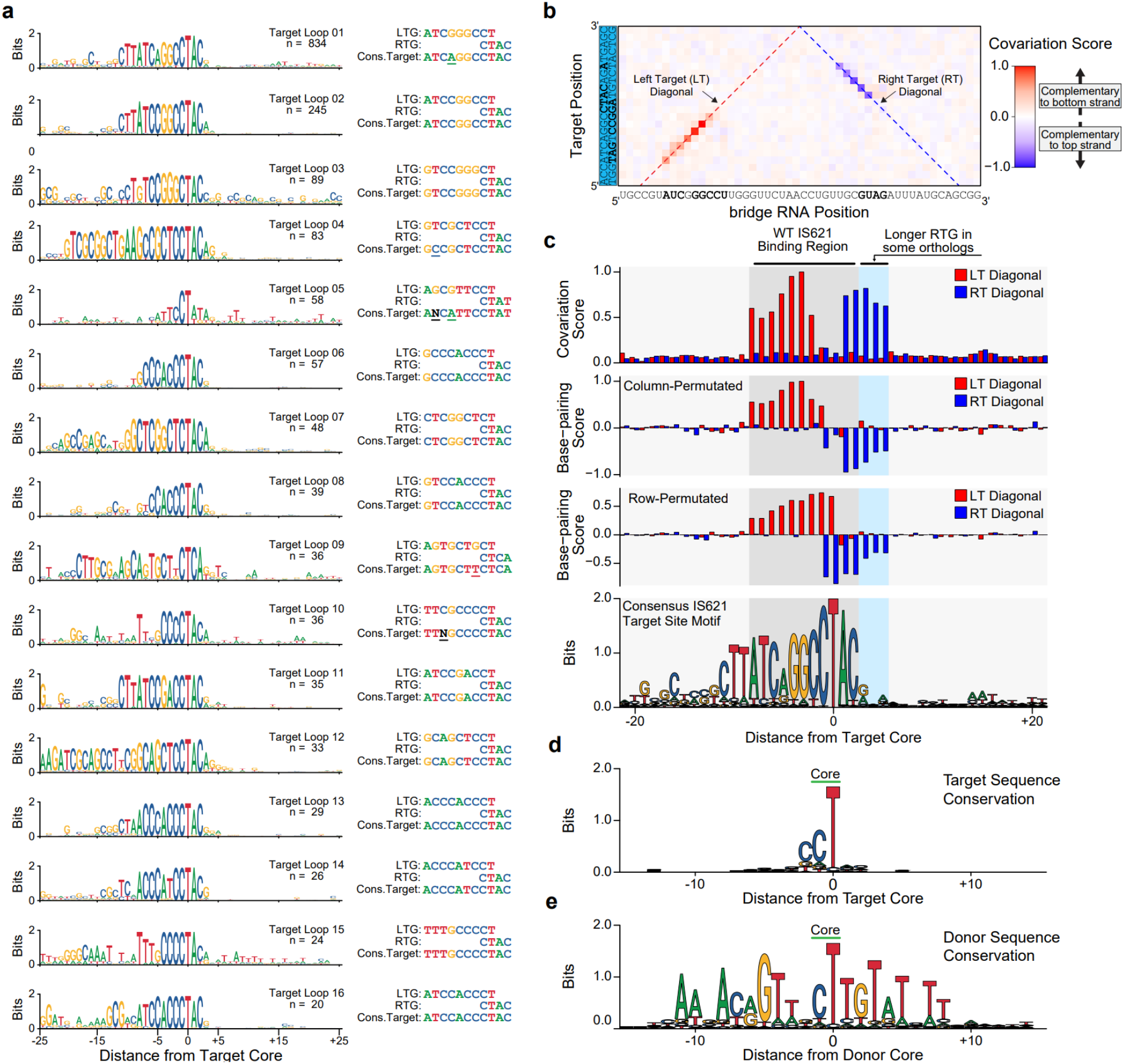
Further details of IS110 covariation and sequence analysis. (**a**) Target sequence motifs associated with 16 distinct bridge RNA target-binding loop guide sequences. All unique guides with 20 or more associated target sequences were retained. For each guide, an 11 bp consensus target sequence was constructed by taking the most abundant nucleotide at each position. If two sequences tied as the most abundant nucleotide, they were represented using ambiguous IUPAC code N. Highlighting the mismatching positions between the target guide and the consensus target. (**b**) Schematic of the target-bridge RNA covariation analysis presented in Fig. 2b with annotations to indicate the left target (LT) diagonal and the right target (RT) diagonal. The values of different metrics along these diagonals are shown in (**c**). (**c**) Boundaries of the programmable positions in the target sequence. The top panel indicates the covariation scores along the LT and RT diagonals as generated by CCMpred, which are normalized between 0 and 1. The second panel shows the column-permuted base-pairing score, which is an additional statistic that can be used to identify nucleotide covariation signals while considering both top-and bottom-strand base-pairing. The sign of this score (+1/-1) is multiplied by the covariation score to generate the covariation signals shown in **Fig. 2b** and **Supplementary Fig. 4**. The third panel shows the row-permuted base-pairing score. The bottom panel shows a sequence logo for all identified IS621 insertion sites. All these panels are aligned with respect to the core of the target sequence. (**d**) A sequence logo of 5,485 diverse IS621 target sequences. All unique target and donor sequences that were used as input in the covariation analysis are shown here. (**e**) A sequence logo of 5,485 diverse IS621 donor sequences. The cognate donor sequences for the targets shown in (**e**) are represented here.

**Supplementary Figure 6:**
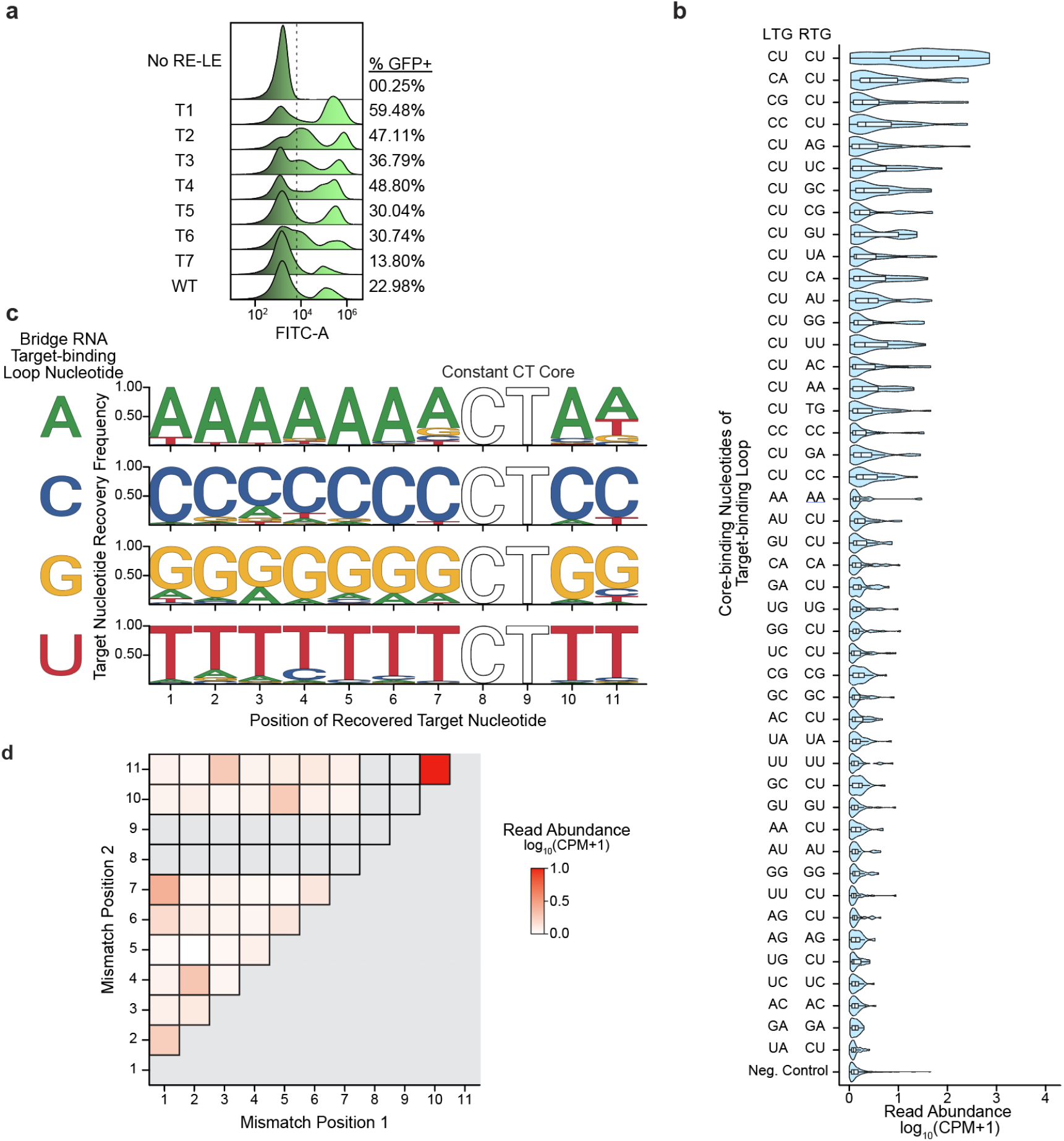
Extended results of RNA target-binding loop and target reprogramming. (**a**) DNA recombination in *E.* coli with reprogrammed bridge RNAs. The distribution of FITC-A signal for the cell population is shown, with a representative gating strategy for evaluating the percentage of GFP+ cells. Plots are representative of 3 replicates featured in Figure 3d. (**b**) Read abundance of oligos with bridge RNA target-binding loop mutations at the positions that bind to the core sequence. The 2 base-pair nucleotides predicted to bind the core in the target-binding loop LTG and RTG were mutated while holding the target and donor CT cores constant and varying the 9 other programmable positions. All tested core mutation combinations shown here, along with a negative control set of 9 mismatch target/target-loop combinations. (**c**) Mismatch tolerance at each position of the 11 bp target sequence. The x-axis shows the target position, with the CT core held constant. The top panel shows the target nucleotide recovery frequency when the target-binding loop contains an A at each guide position, the second panel shows the same but when the target-binding loop contains a C at each position, etc. as a percentage of recovered recombinants at each position. (**d**) Double mismatch tolerance for combinations of positions within the target and target-binding loop. Each cell indicates the average read abundance of oligos that contain double mismatches at the two corresponding positions. The core was held constant.

**Supplementary Figure 7:**
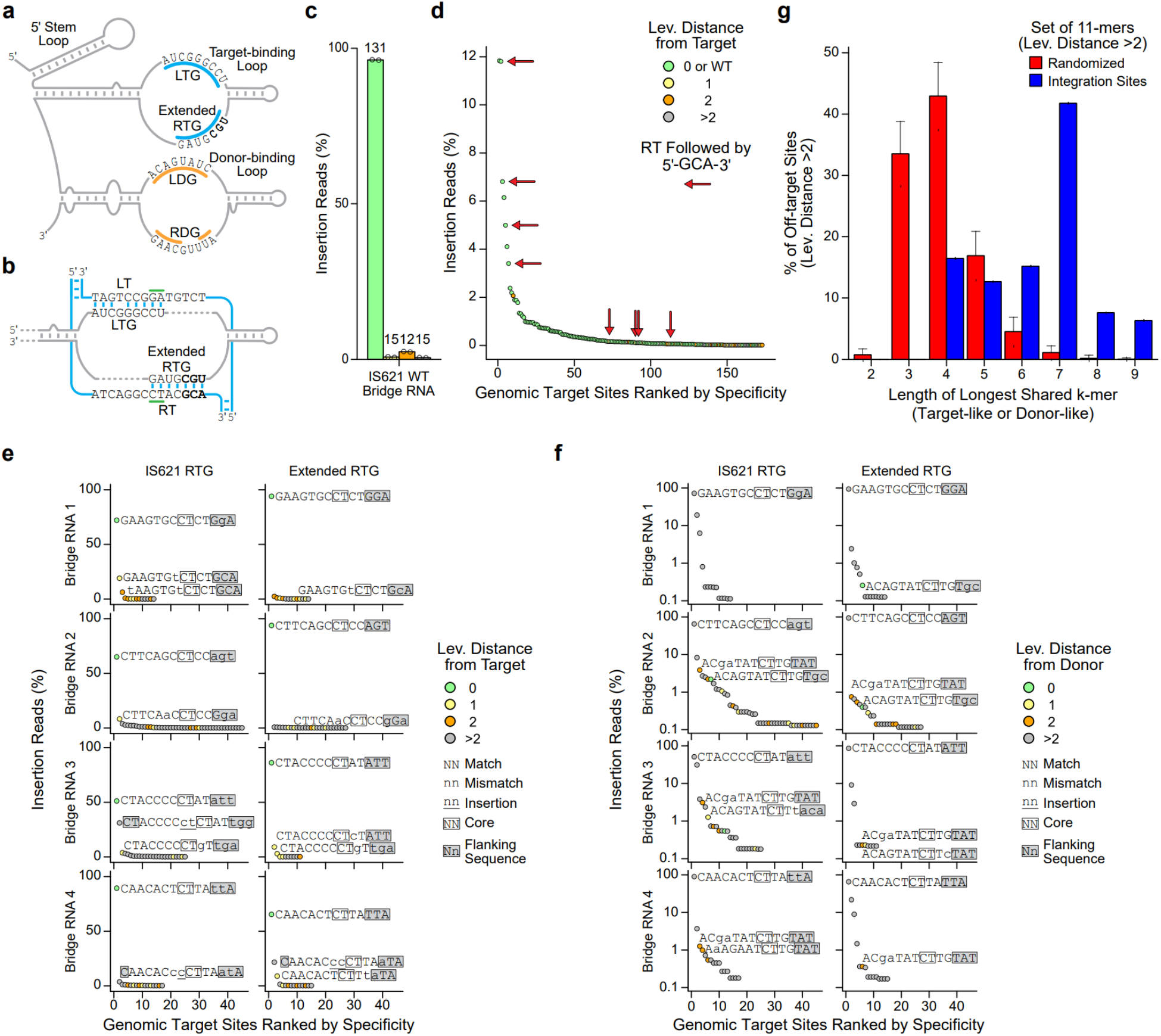
Effect of extended RTGs on specificity of genome integration by IS621 bridge RNA and recombinase. (**a**) Schematic indicating the bases of the WT bridge RNA which may represent an extended RTG. (**b**) Schematic indicating how the target-binding loop of the WT bridge RNA can form more base pairs with a WT 11 bp target sequence flanked on the 5′ end by 5′-GCA-3′. (**c**) Genomic specificity profile of the IS621 WT bridge RNAs. Color indicates the number of differences from the intended sites as measured by Levenshtein distance. Data represent sums of all insertion sites with 0 or WT (ATC<u>A</u>GGCCTAC), 1, 2 or >2 differences from the expected target. (**d**) Insertion sites into the *E. coli* genome ranked by abundance. Insertion sites where the RTG flanking bases match the RT flanking bases are indicated with red arrows. (**e**) Genomic insertion sites for reprogrammed bridge RNAs displayed in rank order. High frequency insertion sites are highlighted by descriptions of similarity to the intended target sequence. Color indicates the number of differences from the expected sites as measured by Levenshtein distance. (**f**) Genomic insertion sites in (**e**) recolored to represent similarity to the WT donor sequence. (**g**) Similarity of off-target sites (Lev. Distance > 2 from expected target and donor) to the expected target and donor sequences as measured by the length of the longest shared *k*-mer. For comparison, off-target sites were randomly shuffled and the procedure was repeated 1,000 times to generate a null distribution. Error bars indicate the standard deviation of the 1,000 permutations.

**Supplementary Figure 8:**
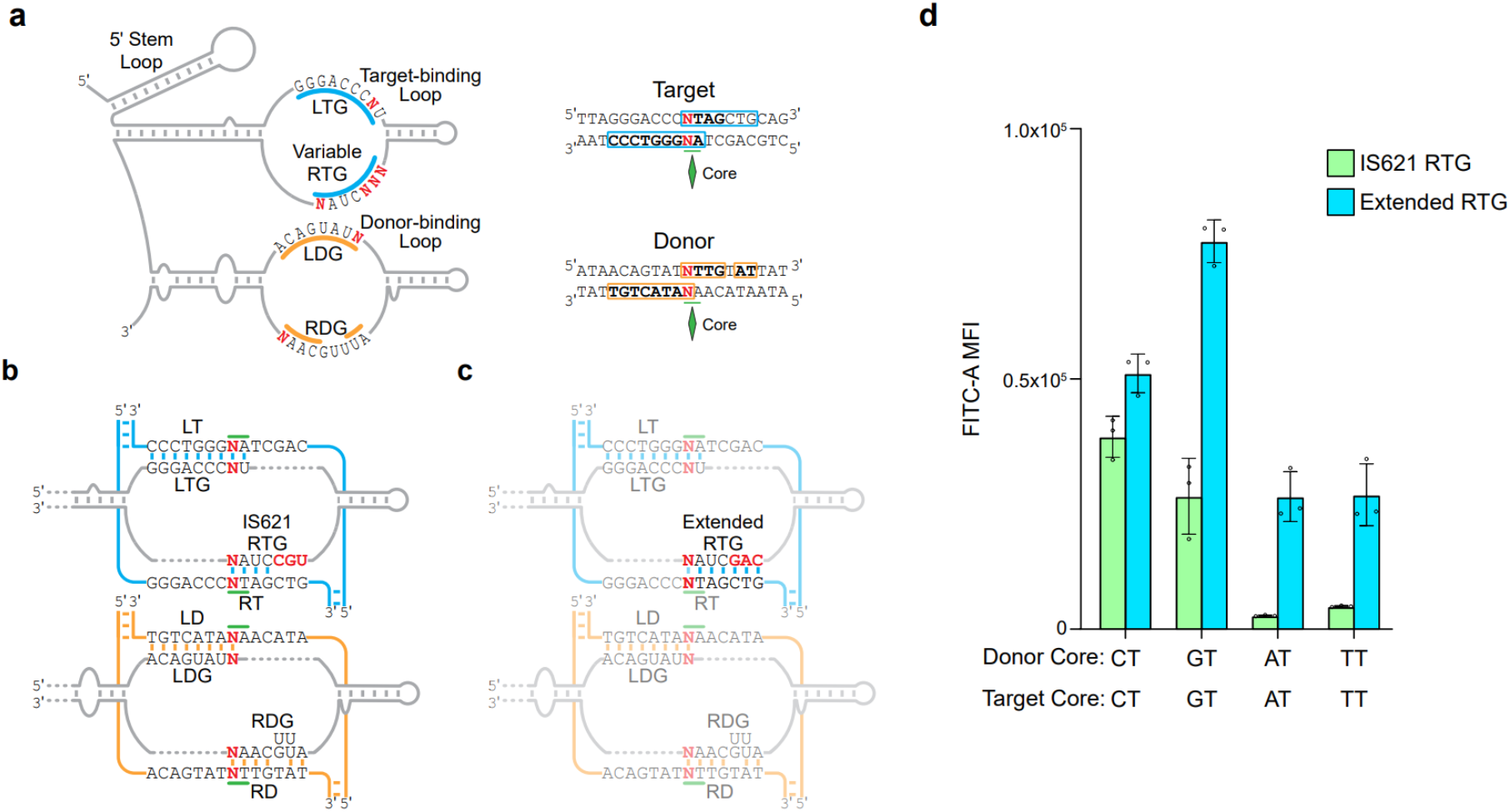
Core reprogramming is enhanced by using a 7 nucleotide RTG. (**a**) Schematic representation of core reprogramming with or without RTG extension. The four positions in the guide loops which bind the first base of the core were mutated along with the first base of the core in the target/donor sequences to test if the core sequence can be reprogrammed. The RTG was programmed to allow 7 base binding with the RT or kept as the WT. (**b**) Schematic representation of base pairing between a target-binding loop with the WT IS621 RTG. (**c**) Schematic representation of base pairing between a target-binding loop with a reprogrammed and extended RTG. (**d**) Plasmid recombination GFP reporter assay to assess the impact of extended RTG on core reprogrammability. The canonical CT core was mutated to GT, AT, and TT, and tested with the IS621 WT RTG (4 bp) and with an extended RTG (7 bp). MFI ± SD for three biological replicates shown.

**Supplementary Figure 9:**
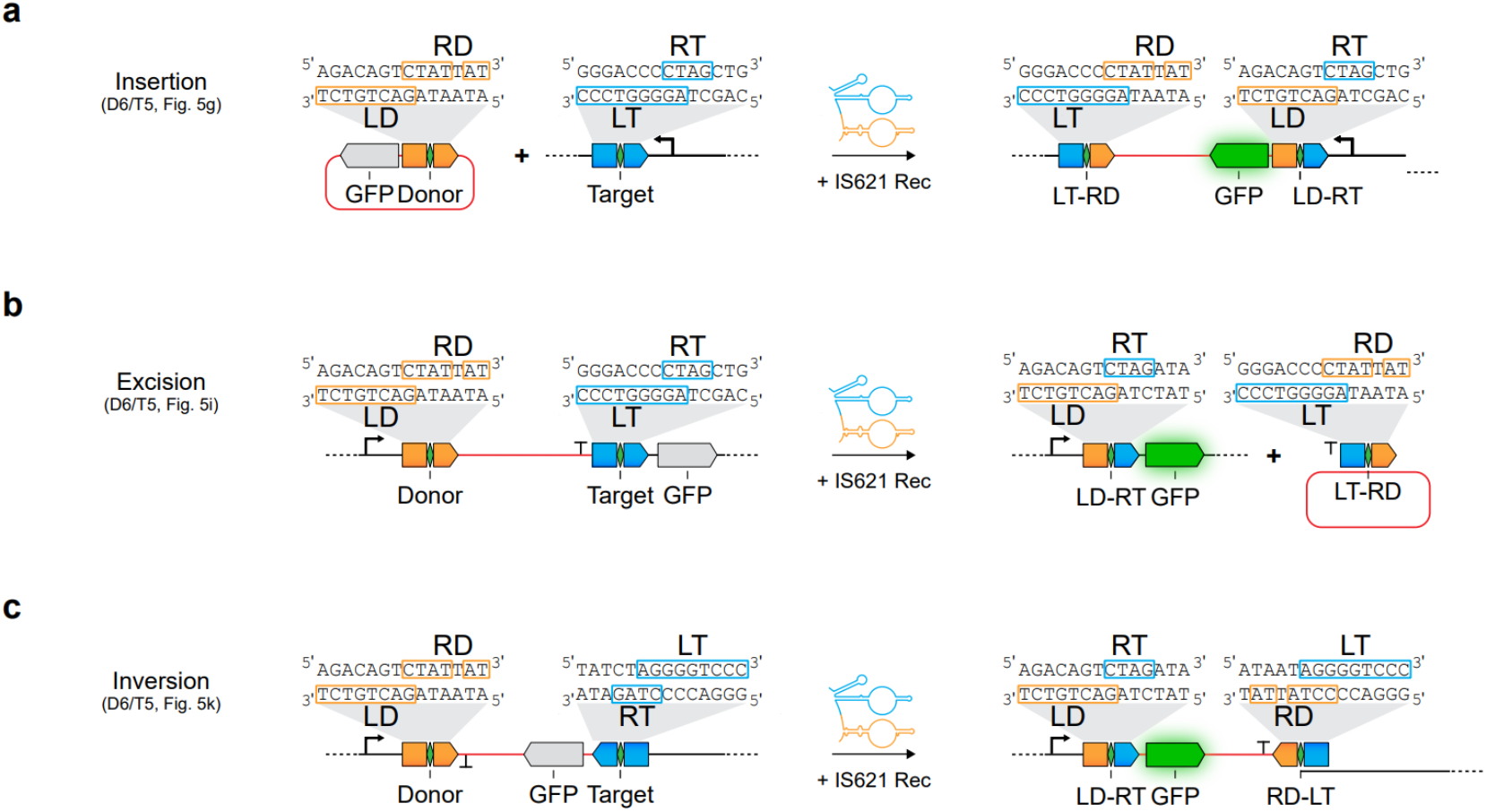
Schematic representations of insertion, excision, and inversion reactions. (**a**) Schematic of the insertion reaction. Insertion takes place when the target and donor sequences are on different DNA molecules. The orientation of the insertion can be controlled by the strand placement of the target and the donor. (**b**) Schematic of the excision reaction. Excision can occur when the target and donor sequences exist on the same molecule and in the same orientation (i.e. LD and LT are on the same strand). (**c**) Schematic of the inversion reaction. Inversion can be catalyzed when the target and donor sequences exist on the same molecule, but in the opposing orientation (i.e. on opposite strands).

**Supplementary Figure 10:**
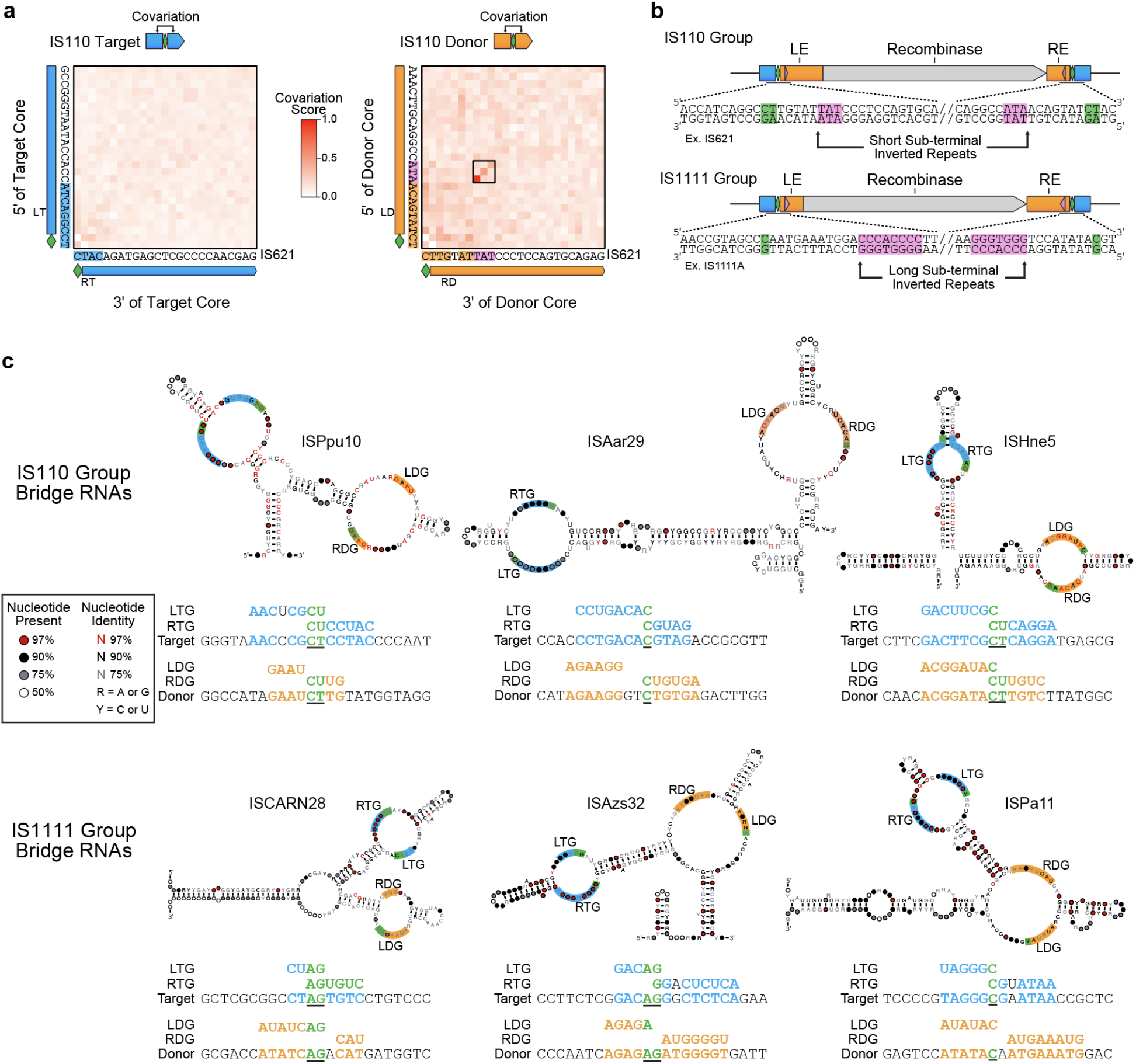
Detailed analysis of diverse bridge RNA sequences and their predicted target/donor binding patterns. (**a**) Covariation analysis of IS110 donor sequences identifies a short sub-terminal inverted repeat (STIR). Target and donor sequences were analyzed using the same covariation analysis introduced in **Fig. 2b**. Target sequences have no notable covariation signal while donor sequences have a prominent 3-base covariation signal that corresponds with an LT-flanking ATA tri-nucleotide and a RD-flanking TAT tri-nucleotide. (**b**) Schematic depicting sequence features of IS110 and IS1111 group elements. IS110 are characterized by long LEs, short REs, and short STIRs. IS1111 are characterized by short LEs, long REs, and long STIRs. (**c**) Six diverse bridge RNAs and their predicted binding patterns. The bridge RNA consensus structures shown are the same as those presented in **Fig. 6d**, but with more detail. Secondary structures are shown with internal loops colored according to the sequence that they complement - target (blue), donor (orange), or core (green). Three members of each IS110 group are shown. For each of the 6 sequence elements cataloged ISfinder - ISPpu10, ISAar29, ISHne5, ISCARN28, ISAzs32, and ISPa11 - IS element boundaries were inspected to identify possible base-pairing between the loops, the targets, and the donors. Under each structure, the predicted LTG, RTG, target, LDG, RDG, and donor are all shown and aligned with respect to the core (underlined in black).

## METHODS

### Development of metagenomic and genomic sequence database

A custom sequence database of bacterial isolate and metagenomic sequences was constructed by aggregating publicly available sequence databases, including NCBI, UHGG (Almeida et al. 2021), JGI IMG (Chen et al. 2021), the Gut Phage Database (Camarillo- Guerrero et al. 2021), the Human Gastrointestinal Bacteria Genome Collection (Forster et al. 2019), MGnify (Mitchell et al. 2020), Youngblut et al animal gut metagenomes (Youngblut et al. 2020), MGRAST (Meyer et al. 2008), and Tara Oceans samples (Sunagawa et al. 2015). The final sequence database included 37,067 metagenomes, 274,880 bacterial and archaeal metagenome-assembled genomes (MAGs), 855,228 bacterial and archaeal isolate genomes, and 185,140 predicted viral genomes.

### Analysis of conserved residues in IS110 protein sequences

Genomic sequences were annotated using Prodigal (Hyatt et al., 2010) to identify coding sequences (CDS). All unique protein sequences were then combined into a single FASTA file and clustered at 30% sequence identity using mmseqs2 (Steinegger & Söding, 2017). Two Pfam domains DEDD_Tnp_IS110 (PF01548) and Transposase_20 (PF02371) were used to search against these clustered representative proteins using the hmmsearch tool in the hmmer package (Finn et al., 2011). DEDD_Tnp_IS110 was used to identify the RuvC-like domain, and Transposase_20 was used to identify the Tnp domain. All members of the matched 30% identity clusters were then extracted, and the same IS110 Pfam domain significance thresholds were applied to filter these candidates. Next, only proteins that met E < 1e-3 for both domains were retained. Next, RuvC-like domains were only retained if they were between 125 and 175 aa in length, and Tnp domains were only retained if they were between 60 and 110 aa in length. Any sequences with ambiguous residues were removed. Protein domains were then clustered at 90% using mmseqs (“easy-cluster --cluster-reassign -c 0.8 --min-seq-id 0.9 --cov-mode 0”). Cluster representatives were then aligned using hmmalign (“--trim --amino”) (Finn et al., 2011). Alignment columns with more than 50% gaps were removed, and the alignments were visualized using ggseqlogo in R (Wagih, 2017).

### Phylogenetic analysis of IS110 transposases

A phylogenetic analysis of IS110 transposases was also performed. Full-length IS110 proteins were clustered at 90% identity using the mmseqs2 easy-cluster algorithm (“--cluster-reassign -c 0.85 --min-seq-id 0.9 --cov-mode 0”) (Steinegger & Söding, 2017) . Next, using the identified 90% protein sequence clusters, a representative from each cluster was selected that was closest to the 80th percentile in total length. This resulted in a curated set of 90% identity cluster representatives. Next, 90% identity cluster representatives were clustered at 30% identity across 70% of the aligned sequences using the mmseqs2 easy-cluster algorithm (“--cluster-reassign -c 0.70 --min-seq-id 0.30 --threads 96 --cov-mode 0”). This resulted in 1,686 30% identity cluster representatives. RuvC-like and Tnp-like domains were extracted from these proteins using the corresponding Pfam pHMM models and hmmsearch (Finn et al., 2011). These extracted domains were then individually aligned using hmmalign (“--amino --trim”) and concatenated into a paired alignment. All pairwise percent identity values were calculated for this alignment, and redundant sequences were removed using a 60% identity cutoff, resulting in 1,054 aligned sequences. A phylogenetic tree was then constructed using iqtree2 v2.1.4-beta, with all default parameters (Minh et al., 2020), midpoint-rooted, and visualized in R with ggtree (Yu, 2020). Additional metadata about each sequence was mapped onto the tree, including host kingdom and phylum, ISfinder group, and notable orthologs.

Curated ISfinder transposases were analyzed separately to produce another phylogenetic tree. IS110 transposase sequences were extracted from the database available through the prokka software package (Seemann, 2014). Only IS110 transposases over 250 aa were retained. Protein sequences were then clustered using mmseqs2 (“easy-cluster -c 0.5 --min-seq-id 0.9 --threads 8 --cov-mode 0”) (Steinegger & Söding, 2017). Cluster representatives were then aligned using mafft-ginsi (“--maxiterate 1000”) (Katoh et al., 2002). Alignment columns with over 50% gaps were removed. A phylogenetic tree was then constructed using iqtree2 v2.1.4-beta with all default parameters (Minh et al., 2020).

### Analysis of LE and RE lengths across IS110 elements

Sequence coordinate information about individual IS elements was collected through the ISfinder web portal (Siguier et al. 2006). This included information about the total length of each IS element, as well as the start and end coordinates of the recombinase CDS. The LE non-coding length was calculated from the CDS coordinates for each IS110 element as the distance between the 5′ terminus and start of the CDS, and the RE non-coding length was calculated as the distance between the end of the CDS and the 3′ terminus. Tn3 family elements were excluded due to highly variable passenger gene content.

### Predicting IS110 element boundaries

To identify the boundaries of each element, an initial search was conducted using comparative genomics to identify putative pre-insertion and post-insertion examples within the metagenomic sequence database. IS110 protein candidates were clustered at 30% identity using mmseqs2 (Steinegger and Söding 2017), and within each cluster all relevant genomic loci were identified. Nucleotide sequences were then extracted from the database by adding 1,000 base pairs to the 5′ and 3′ ends of the IS110 CDS, and extracting the complete intervening sequence. These IS110 loci were then separated into “batches” based on 90% identity protein clusters. These batches were then searched against up to 40 metagenomic or isolate samples in the custom database, prioritizing samples that already contained related recombinases. Putative pre-insertion sites were identified if the distal ends of the loci aligned by BLAST to a contiguous sequence (Altschul et al. 1990), but the IS110 CDS did not. Precise boundaries of the IS110 element were then predicted using a modified method similar to what was implemented by the previously published tool MGEfinder (Durrant et al., 2020). Core sequences were identified as repeated sequences near the end of the predicted element. Next, an iterative BLAST search was used to extend IS110 element boundary predictions beyond those that could be detected by identifying pre-insertion sites. IS110 elements were searched using BLAST against all IS110 loci. Hits were retained only if both ends of the element aligned, and if the core was concordant between query and target. This then generated a new set of IS110 elements and their boundaries, which were recycled as query sequences, and the search was repeated for another iteration. This repeated for 36 iterations before convergence (no new IS110 elements were found). The combined set of IS110 boundaries were kept for further analysis.

### Identification of bridge RNA consensus structures

A pipeline was developed to identify conserved RNA structures in the sequences immediately flanking the recombinase CDS. First, the IS621 protein sequence was searched against the complete IS110 database for orthologs using blastp (“-max_target_seqs 1000000 -evalue 1e-6”). Only hits that were at least 30% identical at the amino acid level with 80% of both sequences covered by the alignment were retained. Up to 2000 unique proteins were then selected in order of descending percent amino acid identity. Flanking sequences for the corresponding proteins were then retrieved from the database, with flanking sequences defined as a 5′ flank of up to 255 bp (including 50 bp of 5′ CDS) and a 3′ flank of up to 170 bp (including 50 bp of the 3′ CDS). These flanks were then further filtered to exclude sequences that were more than 35 bases shorter than the target flank lengths. Sequences were filtered to exclude those with ambiguous nucleotides. Protein sequences were then clustered using mmseqs2 easy-linclust with a minimum percent nucleotide identity cutoff of 95% across 80% of the aligned sequences, and one set of flanks for each representative was retained. Flanking sequences were then clustered at 90% nucleotide identity across 80% of the aligned sequences, and only one representative flanking sequence pair per cluster was retained. Then, up to 200 sequences were selected in order of decreasing percent identity shared between the IS621 protein sequence and their corresponding ortholog protein sequence. The remaining sequences were then individually analyzed for secondary RNA structures using linearfold (Huang et al., 2019). Sequences were then aligned to each other using the mafft-xinsi (IS621 ortholog sequences) or mafft-qinsi (all other ISfinder elements) alignment algorithms and parameter --maxiterate 1000 (Katoh et al., 2002). Alignment columns with over 50% gaps were removed. Conserved RNA secondary structure was then projected onto the alignment, and manually inspected to nominate bridge RNA boundaries. This region was exported as a separate sequence alignment file, and a consensus RNA secondary structure was predicted using ConsAlifold (Tagashira & Asai, 2022). This structure was then visualized using R2R (Weinberg & Breaker, 2011). This same pipeline was used to analyze hundreds of other IS110 elements, resulting in diverse predicted secondary structures. For visualization purposes, consensus secondary structures with minimally structured terminal ends were trimmed to the primary structured sequence. These consensus structures were converted into covariance models using infernal (Nawrocki & Eddy, 2013), and these were then searched across thousands of sequences to identify putative bridge RNAs (Nawrocki and Eddy 2013).

### Nucleotide covariation analysis to identify bridge RNA guide sequences

To identify programmable guide sequences in the bridge RNA of the IS621 element, the following approach was taken. First, the IS621 protein sequence was searched against our collection of IS110 recombinase proteins with predicted element boundaries using blastp. Next, only alignments that met a cutoff of 20% amino acid identity across 90% of both sequences were retained. Next, a covariance model (CM) of the bridge RNA secondary and primary sequence was used to identify homologs of the bridge RNA sequence in the non-coding ends of these orthologous sequences (Nawrocki and Eddy 2013). 50 nucleotide target and donor sequences were extracted centered around the core. For elements with multiple predicted boundaries, boundaries with a CT dinucleotide core were prioritized. Next, elements that were identified at earlier iterations in our boundary search were prioritized. Next, elements that were similar in length to the known IS621 sequence element were prioritized. Only 1 element per unique locus was retained. Alignments were further filtered to remove redundant examples by clustering target/donors and bridge RNA sequences at 95% identity, taking 1 representative per pair, and then taking at most 20 examples for each 95% identity bridge RNA cluster. Predicted bridge RNA sequences were then aligned using the cmalign tool in the Infernal package (Nawrocki and Eddy 2013). Two paired alignments were then generated that contained concatenated target and bridge RNA sequences, and concatenated donor and bridge RNA sequences. These alignments were then further filtered to remove all columns that contained gaps in the IS621 bridge RNA sequence. These alignments were then analyzed using CCMpred (“-n 100”) to identify co-varying nucleotides between target/donor and bridge RNA sequences (Ekeberg et al., 2013). These covariation scores were normalized by min-max normalization and multiplied by the sign of the column-permuted base-pairing concordance score (see next paragraph), with +1 corresponding with bottom strand base-pairing and -1 corresponding with top strand base-pairing. The signal was visualized as a heat map and interactions were identified within the two internal loops of the bridge RNA, leading to the proposed model for bridge RNA target/donor recognition. The same covariation analysis was performed on the donor alone, leading to the identification of short STIR sequences for IS110 elements.

A separate analysis was performed on the same paired alignment used in the covariation analysis to determine if certain pairs of nucleotides were biased toward base-pairing. The observed concordance was first calculated for each pair of columns as:

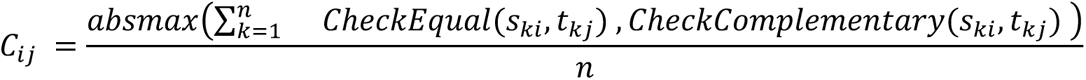

Where 𝐶 is the concordance score, 𝑖 refers to the first column (or position), 𝑗 refers to the second column, 𝑛 refers to the total number of rows (sequences) in the alignment, 𝑠*_ki_* refers to the nucleotide in bridge RNA sequence 𝑘 at position 𝑖, and 𝑡*_kj_* refers to the nucleotide in target (or donor) sequence 𝑘 at position 𝑗. 𝑎𝑏𝑠𝑚𝑎𝑥(𝑎, 𝑏) is a function that returns the value with the largest absolute magnitude, 𝐶ℎ𝑒𝑐𝑘𝐸𝑞𝑢𝑎𝑙(𝑎, 𝑏) is a function that returns one when 𝑎 = 𝑏 and 0 otherwise, and 𝐶ℎ𝑒𝑐𝑘𝐶𝑜𝑚𝑝𝑙𝑒𝑚𝑒𝑛𝑡𝑎𝑟𝑦(𝑎, 𝑏) is a function that returns -1 if 𝑎 and 𝑏 are complementary nucleotides and 0 otherwise. All positions where the nucleotide is a gap in either sequence are ignored and discounted from 𝑛. All observed values of 𝐶*_ij_* are then compared with two different null distributions of 𝐶*_ij_* scores. The first is generated by randomly permuting the rows of the bridge RNA alignment 1000 times and recalculating 𝐶 for each permutation, and the second is generated by randomly permuting the columns of the bridge RNA alignment 1000 times and recalculating 𝐶. The mean and standard deviation of these permuted 𝐶 distributions is then used to convert the observed 𝐶 scores into Z-scores, and positive and negative values are then separately min-max normalized to maintain the -1 to 1 scale. The sign of this score is then used to project base-pairing information onto the covariation scores as generated by CCMpred.

### Small RNA sequencing of IS110 bridge RNAs

BL21(DE3) *E. coli* were transformed with plasmids bearing a concatenated RE-LE sequence and plated on an LB agar plate with appropriate antibiotics. A single colony was picked and grown in terrific broth (TB) to an optical density (OD) of 0.5.RNA isolation was performed using Direct-zol RNA Miniprep Kit (Zymo Research). RNA was prepared for small RNA sequencing according to the following protocol. Briefly, no more than 5µg total RNA was treated with DNase I (NEB) for 30 minutes at 37°C then purified using RNA Clean & Concentrator -5 Kit. Ribosomal RNA was depleted from samples using Ribo-Zero Plus rRNA Depletion Kit (Illumina) and purified using RNA Clean & Concentrator - 5 Kit. Depleted RNA was treated with T4 PNK for six hours at 37°C, supplementing with T4 PNK and ATP after six hours for one additional hour. RNA was purified using RNA Clean & Concentrator - 5 kit and subsequently treated with RNA 5′ Polyphosphatase (Lucigen) for 30 minutes at 37°C. RNA was purified with RNA Clean & Concentrator - 5 Kit and concentration was measured via nanodrop. Next-generation sequencing libraries were prepared using NEBNext Multiplex Small RNA Library Prep Kit (NEB) according to the manufacturer’s protocol. Libraries were sequenced on an Illumina MiSeq using a 2x150 Reagent Kit v2.

### Analysis of small RNA sequencing data

Demultiplexed fastq files were cleaned and merged using bbduk and bbmerge, respectively (Bushnell, Rood, and Singer 2017). Merged fastq files were aligned to the RE-LE bearing plasmid using bwa mem (Li and Durbin 2009). Small RNA-seq coverage was normalized according to the maximum read depth observed for each ortholog across the entire RE-LE plasmid.

### In vitro transcription of bridge RNAs

*In vitro* transcription was carried out on a linear DNA template using the HighScribe T7 High Yield RNA Synthesis Kit (New England Biolabs) as per the manufacturer’s instructions. DNA template was prepared by cloning into a pUC19 backbone and plasmid was linearized using SapI restriction enzyme (NEB) and purified using DNA Clean & Concentrate (Zymogen). After IVT, RNA was purified using the Monarch RNA Cleanup Kit. Where necessary, bridge RNA was further purified denaturing polyacrylamide gel electrophoresis, extracted from the gel using UV shadowing and recovered by ethanol precipitation.

### IS621 protein preparation

The IS621 recombinase gene was human codon optimized and cloned into a modified pFastBac expression vector (Addgene, Item ID: 30115), which includes an N-terminal His6-tag, a Twin-Strep-tag, and a Tobacco etch virus (TEV) protease cleavage site. IS621 recombinase protein was expressed in Sf9 cells (ATCC, CRL-1711) and purified via FPLC. Elution fractions containing recombinant protein were concentrated using a 10 kDa MWCO ultracentrifugal concentrator (MilliporeSigma) and buffer exchanged during centrifugation into size-exclusion chromatography (SEC) buffer. SEC was carried out using a Superdex 200 Increase 10/300 GL column (Cytiva) to further purify the protein, and the peak fractions were collected, concentrated, and stored at −80°C until use.

### Microscale Thermophoresis

Microscale thermophoresis (MST) was carried out using a Monolith NT.115^pico^ Series instrument (NanoTemper technologies). IS621 recombinase was labeled for MST using the RED-MALEIMIDE 2^nd^ Generation cysteine reactive kit (NanoTemper technologies) as per the manufacturer’s instructions. Labeled protein was eluted in a buffer containing 20 mM Tris-HCl, 500 mM NaCl, 5 mM MgCl2, 1 mM DTT, 0.01% Tween20, pH 7.5. In order to determine affinity of recombinase for RNA, 20 nM recombinase were incubated with a dilution series (2500 - 0.076 nM) of purified LE encoded ncRNA or a scrambled RNA of equivalent length. MST was performed at 37°C using premium capillaries (NanoTemper) at 30% LED excitation and medium MST power. Data were analyzed using the NanoTemper MO.affinity analysis software package and raw data were plotted on Prism for visualization. The binding affinity of IS621 RNP for donor and target DNA, as well as donor and target DNA containing scrambled LD-RD and LT-RT regions were determined using the MST tertiary binding function. Single-strand DNA was purchased from IDT (Coralville, USA) and annealed in buffer containing 10 mM Tris pH 8.0, 5 mM MgCl2 and 5 mM KCl. For MST, 20 nM RNP consisting of labeled IS621 recombinase and LE encoded ncRNA were incubated with a dilution series of duplexed donor or target DNA oligonucleotides (10 µM to 0.076 nM). MST was performed at 37°C using premium capillaries (NanoTemper) at medium MST power with the LED excitation-power set to automatic (excitation ranged from 20-50%).

### *In vitro* recombination assay

*In vitro* activity of IS621 recombinase was evaluated by incubating 10 µM IS621 with 20 µM LE encoded ncRNA and 0.5 µM duplexed target and donor DNA oligonucleotides (Supplementary

Information) in buffer containing 20 mM Tris-HCl, 300 mM NaCl, 5 mM MgCl2, 1 mM DTT, 0.05 U/µL SUPERase•In RNase Inhibitor (Invitrogen) at 37°C for 2 hours. Reactions were then treated with 40 µg Monarch RNaseA (NEB) for 1 hour and then treated with 1.6 units of Proteinase K (NEB) for a further hour before cleanup of DNA with AMPure XP Beads (Beckman Coulter) using a 2X bead ratio. To detect recombination products, 0.5 µL of the purified reaction product was PCR amplified with primers designed to amplify the LT-RD and LD-RT recombination products. PCR products were visualized by running PCR reactions on 8% TBE gel (Invitrogen) and staining with SYBR Safe (Thermo Fisher Scientific) and imaged on a ChemiDoc XRS+ (Bio-Rad). PCR products were sequenced using Oxford Nanopore sequencing (Primordium Labs).

### GFP reporter plasmid-plasmid recombination assay in *E. coli*

BL21(DE3) cells (NEB) were co-transformed with a pTarget plasmid encoding a target sequence and a T7-inducible IS621 recombinase and a pDonor plasmid encoding a bridge RNA, a donor sequence, and GFP CDS upstream such that upon recombination into pRecombinant GFP expression would be activated by the synthetic Bba_R0040 promoter adjacent to the target site. When expressing the bridge RNA *in cis*, pDonor encodes a full length RE-LE sequence (298 bp), which naturally encodes the donor, the bridge RNA, and a promoter to express the bridge RNA. When expressing the bridge RNA *in trans*, pDonor encodes a shortened donor sequence (22 bp) and a bridge RNA driven by the J23119 promoter and followed by the HDV ribozyme.

To measure excision, a Bba_R0040 promoter is separated from the GFP CDS by the donor site, 1kb of intervening DNA sequence including an ECK120029600 to terminate transcription, and a target site on the same strand. Co-expression of a second plasmid encoding a bridge RNA and a T7-inducible IS621 recombinase results in excision of the intervening 1 kb sequence, yielding GFP expression.

To measure inversion, a Bba_R0040 promoter is encoded adjacent to a top strand donor sequence, followed by a GFP CDS and target sequence encoded on the bottom strand. Co-expression of a second plasmid encoding a bridge RNA and a T7-inducible IS621 recombinase results in inversion of the GFP coding sequence (∼900 bp), yielding GFP expression.

In all GFP reporter assays, co-transformed cells were plated on fresh LB agar containing kanamycin, chloramphenicol, and 0.07 mM IPTG to induce recombinase expression. Plates were incubated at 37°C for 16 hours and subsequently incubated at room temperature for 8 hours. Hundreds of colonies were subsequently scraped from the plate, resuspended in TB, and diluted to an appropriate concentration for flow cytometry. ∼50,000 cells were analyzed on a Novocyte Quanteon Flow Cytometer to assess the fluorescence intensity of GFP-expressing cells. The mean fluorescence intensity of the population (including both GFP+ and GFP-cells) is plotted. pRecombinant plasmids were isolated by picking GFP+ colonies under blue light, seeding in TB containing kanamycin and chloramphenicol, incubating for 16 hours at 37°C shaking at 200 rpm, and isolating using QIAprep Spin Miniprep Kit. The isolated plasmids were sent for whole plasmid sequencing to confirm recombination (Primordium Labs).

### Design of oligo pool for systematic pairwise screen of bridge RNA target-binding loops and targets

A pooled screen was designed to test target and target-binding loop mismatch tolerance and relative efficiency across diverse guide sequences. Several categories of oligos were designed to answer different questions. First, 10,656 oligos were designed to test hundreds of different target guides with single mismatch pairs. That is, for a given target, one position in the guide and the corresponding position in the target to generate all 4x4 = 16 combinations of nucleotides. Target guides were selected to reduce genomic off-targets. Next, 3,600 oligos were designed to test different combinations of double mismatches between target-binding loop and target. Next, 2,000 oligos were designed as an internal set of negative controls by ensuring that none of the 9 programmable positions (excluding the CT core) matched in the target-binding loop and target. Next, another 1,800 oligos were designed to test more single mismatch combinations, but did not include all 4x4 combinations in target and target-binding loop. Finally, 1,610 oligos were designed to test how mismatches in the dinucleotide core of the bridge RNA sequences affected recombination efficiency. One unique barcode per amplicon was assigned at random, ensuring that no 2 barcodes were within 2 mismatches of each other. Each oligo encoded a synthetic Bba_R0040 promoter followed by a target sequence, a unique barcode, the J23119 promoter, and the first 104 bases of the bridge RNA, which includes the 5′ stem loop and target-binding loop. The oligos were ordered as a single pooled library from Twist Bioscience.

### Cloning of oligo pool for systematic pairwise screen of bridge RNA target-binding loops and targets

A vector encoding the final 73 bp of the bridge RNA (the WT donor-binding loop) and a T7-inducible IS621 recombinase was digested using BsaI. The oligo library was amplified with primers encoding overhangs compatible with the digested vector for Gibson cloning. Briefly, the library was cloned into the vector via Gibson cloning, and electroporated in Endura DUO electrocompetent cells (Biosearch Technologies). Hundreds of thousands of colonies were isolated for sufficient coverage of the oligo library, and plasmids bearing library members were purified using Nucleobond Xtra Midiprep Kit (Macherey Nagel).

### Recombination assay with the library of bridge RNA target-binding loops and targets

The plasmid library encoding thousands of target and bridge RNA target-binding loop pairs was co-electroporated into E. cloni EXPRESS electrocompetent cells (Biosearch Technologies) along with a donor plasmid and an inactive Kanamycin resistance gene. Recombination between the two plasmids results in expression of the kanamycin resistance gene, allowing cell survival. After co-electroporation and recovery, cells were plated on bioassay dishes with LB agar. One plating condition, serving as the control, was LB agar with chloramphenicol and ampicillin, which maintain the plasmids but do not induce or require recombination. A second condition was LB agar with chloramphenicol, ampicillin, kanamycin, and 0.1 mM IPTG; IPTG induces recombinase expression, prompting recombination, while kanamycin selects for cells that have induced recombination between the the donor and target plasmid. Both conditions were performed in two replicates. Recombination indicates a compatible target-target-binding loop pair within the library.

Hundreds of thousands of colonies were scraped from the bioassay dishes and had plasmid DNA extracted using Nucleobond Xtra Midiprep Kit (Macherey Nagel). After plasmid DNA isolation, samples were prepared for next generation sequencing. For DNA isolated from the control conditions, a PCR was used to amplify the barcodes specifying target and bridge RNA pairs to measure the distribution of barcodes without selecting conditions. For DNA isolated from selection conditions, a PCR was used to amplify the barcodes specifying target and bridge RNA pairs, with one primer priming from the donor plasmid and the other priming from the target plasmid such that only barcodes from recombinant plasmids were measured. The distribution of barcodes from recombinant plasmids was subsequently compared to the distribution of barcodes under control conditions.

### Analysis of target specificity screen

Amplicon sequences were processed using the bbduk tool (Bushnell et al., 2017). Amplicon sequencing data was then aligned to their respective wildtypes using bwa mem, with ambiguous nucleotides at all variable positions (Li & Durbin, 2009). Barcodes were then extracted from the amplicons using custom python scripts. Barcodes were mapped to the designed barcode library, tolerating single mismatches when making assignments. This resulted in a table of barcode counts per biological replicate. Using custom R scripts, the counts were normalized within each replicate using counts per million (CPM), which converts raw barcode counts into barcode counts per million barcodes. CPM values were then averaged across the two biological replicates in each condition. For the recombinant barcodes, CPM values were then corrected by the control barcode CPM values using a simple correction factor for each barcode, calculated by dividing the expected barcode CPM (assuming a uniform distribution) by the observed barcode CPM. These corrected CPM values were subsequently used in many of the individual analyses. Mismatch tolerance was assessed by limiting the analysis to the top quintile of most efficient 4x4 single mismatch sets, where each set was ranked according to the barcode with maximum efficiency then averaging the percentage of total CPM within each set at each position. The motif of enriched nucleotides at each position was generated by determining the nucleotide composition of the top quintile of most efficient target-binding loop/target pairs (without mismatches), and comparing this to the nucleotide composition of the entire set.

### IS621 genomic insertion assay with long-read nanopore sequencing

A plasmid was prepared encoding a donor sequence adjacent to a constitutively expressed kanamycin resistance gene and a temperature-sensitive Rep101 protein. Plasmid replication of this donor plasmid was eliminated in cells upon growth at 37°C, ensuring that cells encode a single copy of the donor plasmid. A cell line was prepared encoding this donor plasmid by transforming BL21(DE3) and making the resultant cell line chemically competent using Mix & Go preparation kit (Zymo). The temperature sensitive donor plasmid was then transformed with a second plasmid encoding a T7-inducible recombinase and a constitutively expressed bridge RNA. The donor-binding loop of the bridge RNA was programmed to recognize the donor sequence within the donor plasmid and the target-binding loop of the bridge RNA was programmed to recognize a target sequence in the BL21(DE3) *E. coli* genome. After transformation, cells were recovered and plated on 10 cm LB agar plates containing 0.02 mM IPTG, chloramphenicol, and kanamycin; integration of the donor plasmid and expression of kanamycin resistance gene from the genome is required for cell survival. The 1000s of resultant colonies, each with an integration of the donor plasmid into the genome, were scraped from the plate. Genomic DNA was extracted from the pool of colonies using Quick DNA Miniprep plus kit (Zymo). Genomic DNA was then cleaned up using AMpure XP (Beckman Coulter) and sequenced using bacterial genome nanopore sequencing to at least 100x genome coverage.

Sequencing data was downsampled to 200x sequencing depth in reprogrammed bridge RNA experiments, and 1400x depth in the WT bridge RNA experiments. To identify long reads containing potential insertion junctions between plasmid donor and *E. coli* genome (NZ_CP053602.1), all individual reads were programmatically scanned for the presence of the terminal 20 nucleotides of the donor sequence, excluding the core. If a 20 base-pair sub-sequence of a read matched the 5′ terminus or 3′ terminus (allowing for up to 2 mismatches), then the read was split and the flanking sequences were written to separate files. These flanking sequences were then mapped back to the plasmid sequences and the *E. coli* genome using minimap2 (Li 2018), and assigned as originating from the plasmid or the *E. coli* genome according to whichever had the higher alignment score. Reads were then assigned to specific insertion junctions in the *E. coli* genome to identify precise insertion sites. Insertion sites that were within 5 base-pairs of each other were merged together using bedtools merge (Quinlan and Hall 2010) and a representative insertion site was selected. For the reprogrammed bridge RNA genome insertion experiments, additional filters were applied to remove low-quality alignments and account for a low rate (<1%) of cross-sample contamination (possibly due to index hopping). Low-quality predicted insertion sites were excluded only if they met certain criteria, either 1) Occuring at <1% total insertion frequency, a Levenshtein distance of > 2nt from the 11nt target and donor, and supported by a large fraction of clipped reads (>25%, indicating low alignment quality), or 2) Occuring at <1% total insertion frequency, a Levenshtein distance of > 2nt from the 11nt target and donor, and matching a high frequency (>1%) and close target match (<3 nt Levenshtein distance) in a different sample (indicating likely index hopping across samples). The total number of reads per site was subsequently used to determine the insertion specificity for each site.

Off-target sites were evaluated by calculating the Levenshtein distance (LD) between the 11 nt off-target and the 11 nt target and donor sequences. Sequences that were LD>2 from target and donor were further evaluated by searching for shared *k-*mer sequences in the 14 nt off-target, the 14 nt expected target, and the 14 nt donor. To determine if the off-target sequences were enriched for shared target/donor *k*-mers, the maximum-length shared *k*-mer distribution was generated and compared to a null distribution where the 14 nt off-target sequences were randomly shuffled. This shuffling procedure was repeated 1,000 times to calculate the null distribution.

A computational pipeline was developed to identify potential structural variants (50 bp or greater in size) that were independent from the donor plasmid. All long-read nanopore sequences were aligned to the BL21(DE3) *E. coli* genome (NZ_CP053602.1) and the pDonor and pHelper plasmid sequences. Reads that aligned to the pDonor or pHelper sequences were then excluded from the *E. coli* genome alignment. These filtered alignments were analyzed using fgsv v0.0.1 (Nils Homer, Pamela Russell, Tim Fennell, John Didion, n.d.). The tool geNomad was used to annotate a structural variant involving a possible prophage element (Camargo et al., 2023).

For the WT bridge RNA, REP elements were also identified and annotated to determine how frequently they were targeted. REP elements were identified by a BLAST search of three different known REP sequences collected from two different studies (Choi et al., 2003; Tobes & Pareja, 2006). These query sequences included TGCCGGATGCGGCGTAAACGCCTTATCCGGCCTAC, GCCTGATGCGCTACGCTTATCAGGCCTACG, and GCCTGATGCGACGCTGGCGCGTCTTATCAGGCCTACG.

### Design of oligo pool for systematic screen of bridge RNA donor-binding loops and donors

A pooled screen was designed to test donor-binding loop programmability, mismatch tolerance, and relative efficiency across diverse guide sequences. Several categories of oligos were designed to answer different questions. Donor sequences were selected to reduce predicted genomic off-targets. First, 13,593 oligos were designed that included complete single mismatch scans across 100 distinct donors, including all position 4x4 = 16 mismatches with the donor at the corresponding position. Next, 5000 completely random donor guides were selected and paired with a perfectly matching donor for the analysis of a high number of diverse donor sequences. Finally, 2,297 oligos to test single mismatch and double mismatch scans of the WT donor sequence and 4 other functional donors were included. Next, 50 negative control oligos were included that ensured that none of the 9 programmable positions (excluding the CT core) matched in the donor-binding loop and donor. Each oligo encoded a partial sequence of the IS621 RE (52 bp 5′ of the CT core), the reprogrammed donor sequence, and a full length LE (191 bp) encoding a bridge RNA as found in the WT system such that expression of the bridge RNA would be mediated by the natural promoter *in cis*. The donor site sequence and donor-binding loop sequence of the bridge RNA were modified in each member according to the description above, while the target-binding loop of the bridge RNA was constant and programmed to recognize target sequence T5, which is orthogonal to the BL21(DE3) *E. coli* genome. The oligo was flanked on both ends with sequences suitable for golden gate cloning into a desired plasmid backbone. All oligos were ordered as a single pooled library from Twist.

### Cloning of oligo pool for screen of bridge RNA donor-binding loops and donors

First, a vector was constructed encoding a kanamycin resistance gene with no promoter on the bottom strand, followed by the first 61 bp of the IS621 RE sequence. This was followed by a BsaI landing pad site for golden gate cloning, an HDV ribozyme sequence, and a unique molecular identifier of length 12. The UMI backbone was pre-digested via BsaI and the oligo library was cloned into the backbone via golden gate cloning after amplification with appropriate primers such that the full length IS621 RE was reconstituted and the LE harboring the bridge RNA was directly adjacent to the HDV ribozyme sequence. The resultant library was electroporated in Endura DUO electrocompetent cells (Biosearch Technologies). Hundreds of thousands of colonies were isolated for sufficient coverage of the oligo library, and plasmids bearing library members were purified using Nucleobond Xtra Midiprep Kit (Macherey Nagel).

### Recombination assay with the library of bridge RNA donor-binding loops and donors

The plasmid library encoding thousands of donor and bridge RNA donor-binding loop pairs was co-electroporated into E. coli EXPRESS electrocompetent cells (Biosearch Technologies) with a target plasmid encoding the T5 target sequence and a T7-inducible IS621 recombinase. Recombination between the two plasmids results in expression of the kanamycin resistance gene, allowing cell survival. After co-electroporation and recovery, cells were plated on bioassay dishes with LB agar. One plating condition, serving as the control, was LB agar with chloramphenicol and ampicillin, which maintain the plasmids but do not induce or require recombination. A second condition was LB agar with chloramphenicol, ampicillin, kanamycin, and 0.07mM IPTG; IPTG induces recombinase expression, prompting recombination, while kanamycin selects for cells that have induced recombination between the the donor and target plasmid. Both conditions were performed in two replicates. Recombination indicates a compatible target/target-binding loop pair within the library.

Hundreds of thousands of colonies were scraped from the bioassay dishes and had plasmid DNA extracted using Nucleobond Xtra Midiprep Kit (Macherey Nagel). After plasmid DNA isolation, samples were prepared for next generation sequencing. For DNA isolated from the control conditions, a PCR was used to amplify the UMI specifying donor and bridge RNA pairs to measure the distribution of UMIs without selecting conditions. For DNA isolated from selection conditions, a PCR was used to amplify the UMIs specifying donor and bridge RNA pairs, with one primer priming from the donor plasmid and the other priming from the target plasmid such that only UMIs from recombinant plasmids were measured. The distribution of UMIs from recombinant plasmids was subsequently compared to the distribution of UMIs under control conditions. UMIs were initially mapped to donor-bridge RNA pairs by amplifying a region of the input donor library such that the information of all variable sites within the full length of the RE-LE were captured in addition to the adjacent UMI.

### Analysis of donor specificity screen

All amplicon sequence data was pre-processed using bbduk to remove adapters. Next, unique molecular identifiers (UMIs) were mapped to their respective oligos. This was done by aligning to the expected amplicon sequence with ambiguous N nucleotides in all the variable positions using bwa mem (Li & Durbin, 2009). UMIs were then determined from the alignments, and combined with the variable LDG and RDG to guarantee the uniqueness of each UMI to each oligo. Next, control and recombinant samples were analyzed in much the same way as the previously described target screen, but UMIs were counted rather than assigned barcodes. Next, UMI counts were converted to CPM, averaged across two biological replicates, and normalized according to the correction factors calculated in the control condition. These CPM values were then analyzed across different oligo categories to assess mismatch tolerance, how distance from the wild-type donor affects efficiency, and what nucleotide sequences were favored/disfavored at each position in the donor.

### Additional analyses of natural IS110 sequences

Natural IS621 target sites were extracted from the genomic sequence database by searching for exact matches to the 1,277 IS621, excluding the core. These target sequences were then clustered using mmseqs2 and the parameters “easy-cluster --cov-mode 0 -c 0.800 --min-seq-id 0.800” (Steinegger & Söding, 2017). This search and clustering identified 272 distinct target sites, which were then analyzed to identify a conserved target motif and compared with the experimental observed IS621 target sequences in the *E. coli* BL21(DE3) genome.

A paired alignment of target sites and bridge RNA sequences was analyzed to determine how the target site motif changed as the guide RNAs were varied. All aligned bridge RNA sequences that lacked gaps in the 9 base LTG and the 4 base RTG were first identified. Next, only LTG and RTG sequences with CT core guides were selected. Next, only target-binding loops with over 20 associated target sites were kept. For each of these unique remaining target-binding loops, a consensus sequence of the motif was constructed by selecting the most common nucleotide at each of the 11 target positions. If there were ties, then the position was represented by the ambiguous IUPAC character N. These consensus target sites were then compared with the expected target sites to determine how closely they matched.

## Supplementary Information

**Supplementary Data 1. Uncropped gel images**

Uncropped gel image for IVR with full controls.

**Supplementary Data 2. Replicate correlation, target-binding loop and target screen**

Read abundance of library members in target-binding loop and target screen. Results are shown for two replicates for the control, amplifying only the barcode region, and recombinant, amplifying the barcode region across the recombination junction. Identity line shown as blue dashed line. r; Pearson correlation coefficient.

**Supplementary Data 3. Replicate correlation, genome integration data**

Read abundance of genome insertion sites. Two replicates are depicted for each condition. Identity line shown as blue dashed line. r; Pearson correlation coefficient.

**Supplementary Data 4. Replicate correlation, donor-binding loop and donor screen**

Read abundance of library members in target-binding loop and target screen. Results are shown for two replicates for the control, amplifying only the barcode region, and recombinant, amplifying the barcode region across the recombination junction. Identity line shown as blue dashed line. r; Pearson correlation coefficient.

**Supplementary Data 5: Example of flow cytometry gating strategy**

**Supplementary Note 1: Analysis of structural variants in genome insertion assay**

## Data Availability

Code accompanying this study will be available at https://github.com/hsulab-arc/BridgeRNA2024. A python package for designing bridge RNA sequences will be available at https://github.com/hsulab-arc/BridgeRNADesigner, which will also be accessible through a web application.

## Acknowledgements

We thank all members of the Hsu lab for helpful input throughout the course of this project. We also thank the Arc Institute Scientific Publications Team for assistance with the manuscript, K. Vicari for scientific illustration consultation, and J. Carozza and Y. Madrona for biochemistry and molecular biology expertise. N.T.P. was partially supported by the NIH Biology and Biotechnology of Cell and Gene Therapy Training Program (T32GM139780). M.G.D., J.S.A. and S.K are supported by funding from the Arc Institute. H.N. is supported by JSPS KAKENHI Grant Numbers 21H05281 and 22H00403; Takeda Medical Research Foundation; and the Inamori Research Institute for Science. P.D.H. is supported by funding from the Arc Institute, Rainwater Foundation, Curci Foundation, Rose Hill Innovators Program, S. Altman, V. and N. Khosla, and anonymous gifts to the Hsu Lab.

## Author contributions

M.G.D., N.T.P., and P.D.H. conceived the study. M.G.D., S.K., and P.D.H. designed the computational strategy. N.T.P., J.S.A., M.G.D., S.K., and P.D.H. designed experiments. M.G.D. and J.M. performed computational analyses. N.T.P, J.J.P., A.R.J., J.S.A., and M.H. performed experiments. M.G.D., N.T.P, J.J.P., A.R.J., J.S.A., H.N., SK., and P.D.H. analyzed and interpreted the data. P.D.H. supervised the research with support from S.K., M.G.D., N.T.P., and P.D.H. wrote the manuscript with important contributions from A.P. and S.K., and input from all other authors.

