## Supplementary Information for "Bridge RNAs direct modular and programmable recombination of target and donor DNA"

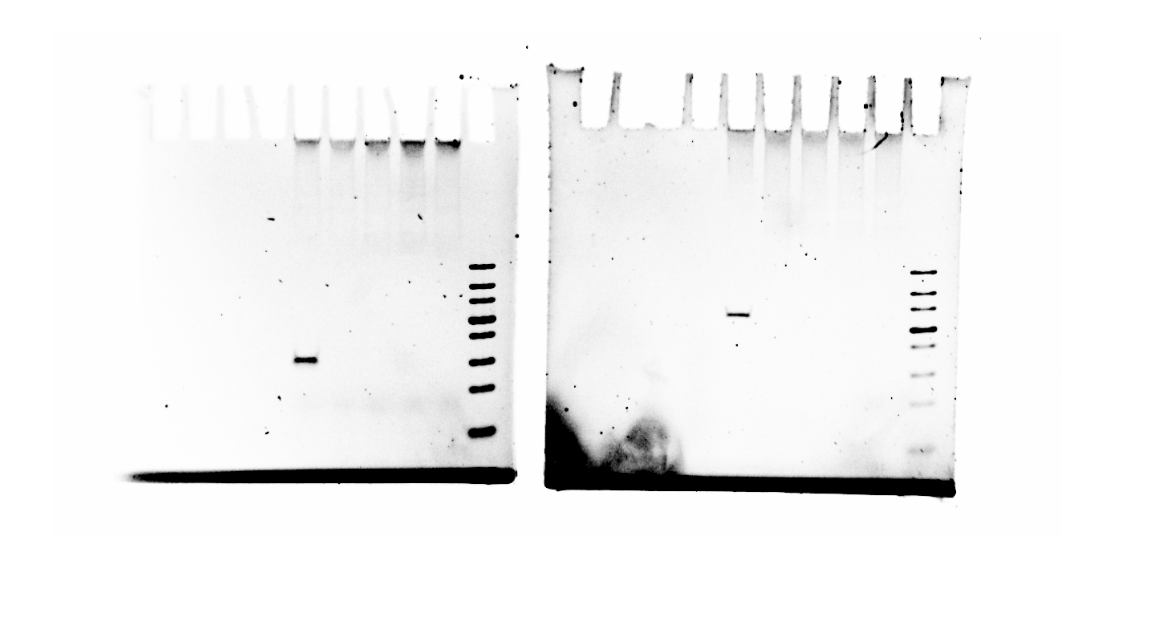

**Supplementary Data 1: Uncropped gel images**

Uncropped gel image for IVR with full controls.

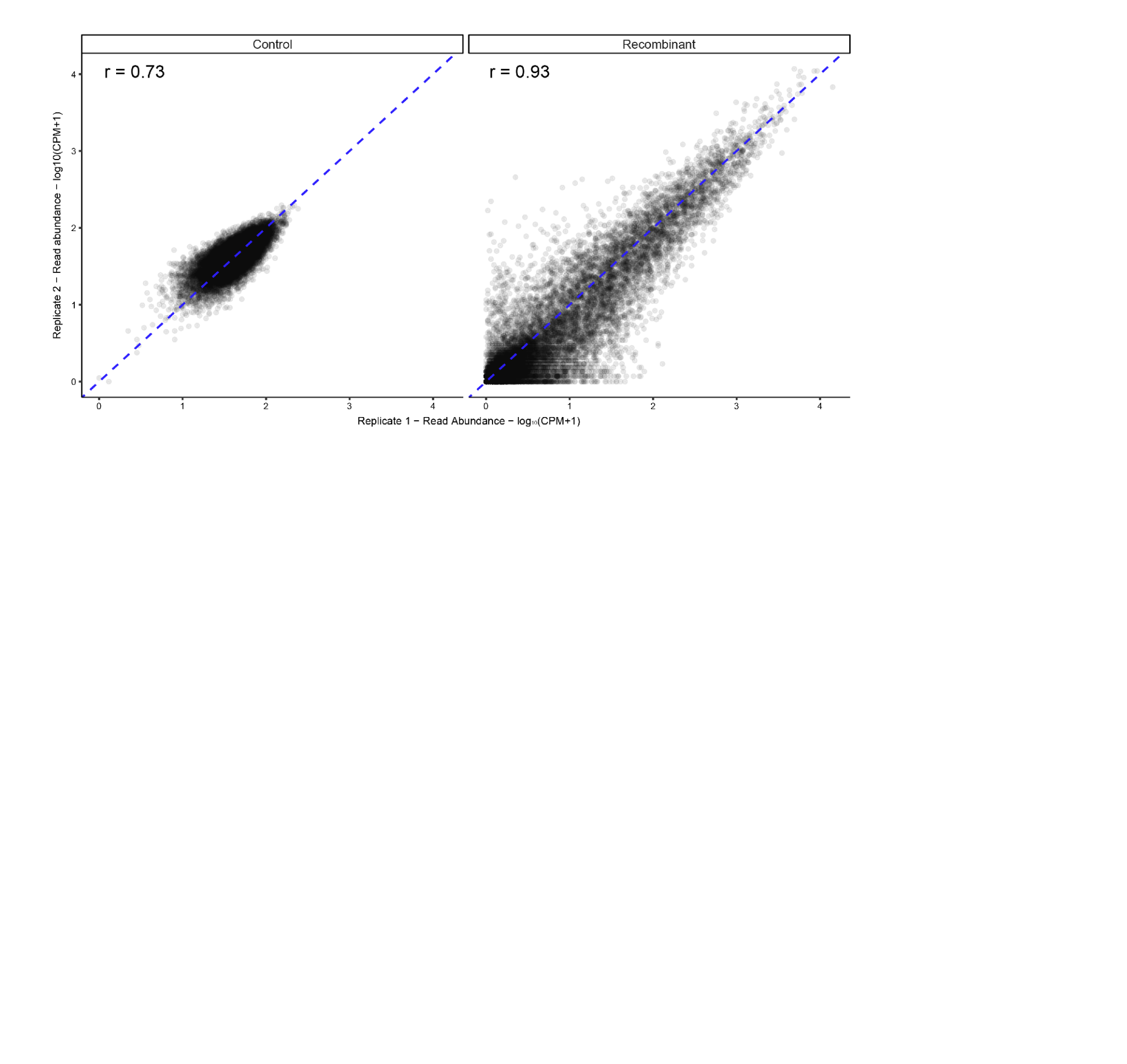

**Supplementary Data 2: Replicate correlation, target-binding loop and target screen**

Read abundance of library members in target-binding loop and target screen. Results are shown for two replicates for the control, amplifying only the barcode region, and recombinant, amplifying the barcode region across the recombination junction. Identity line shown as blue dashed line. r; Pearson correlation coefficient.

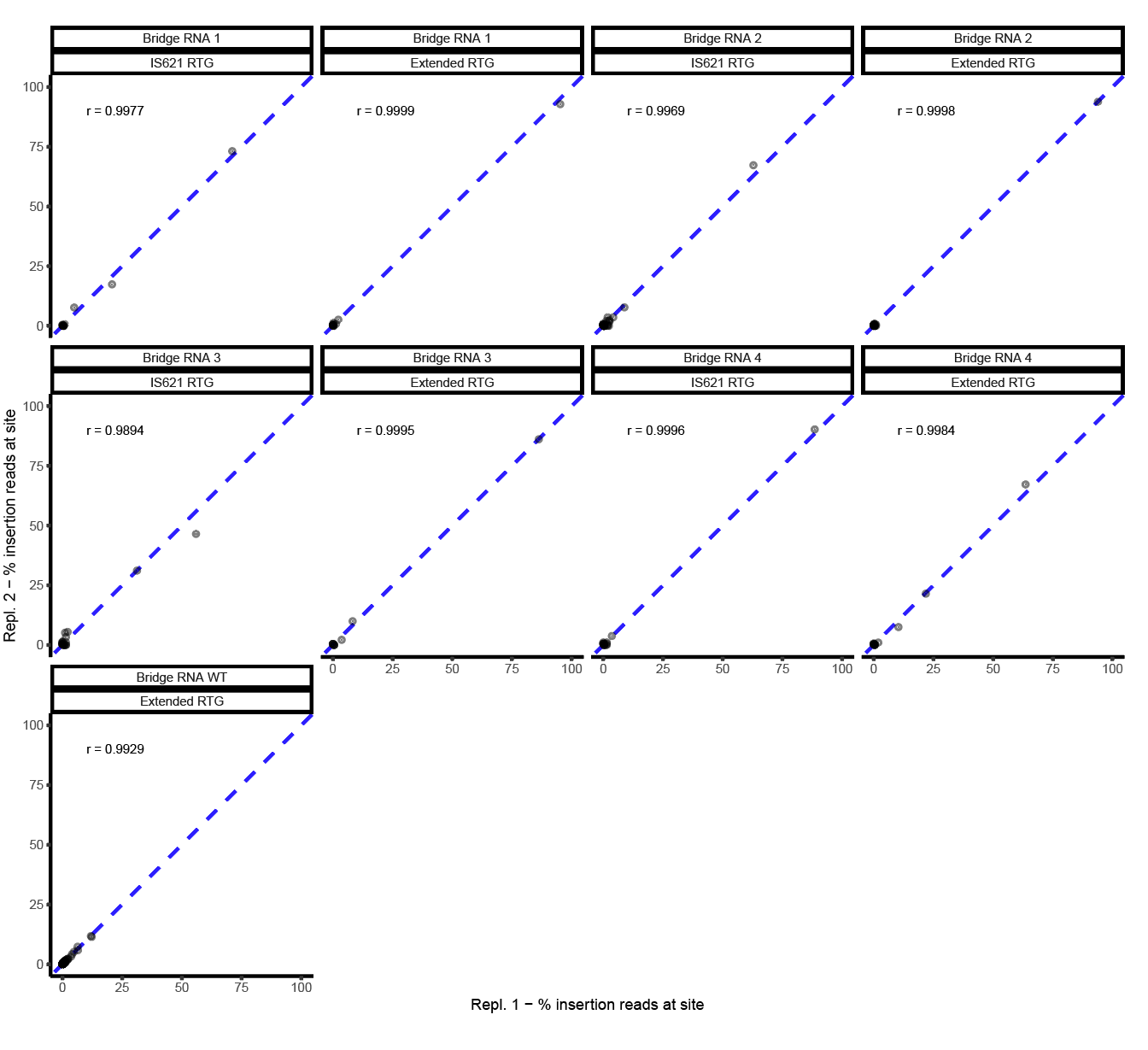

**Supplementary Data 3: Replicate correlation, genome integration data**

Read abundance of genome insertion sites. Two replicates are depicted for each condition. Identity line shown as blue dashed line. r; Pearson correlation coefficient.

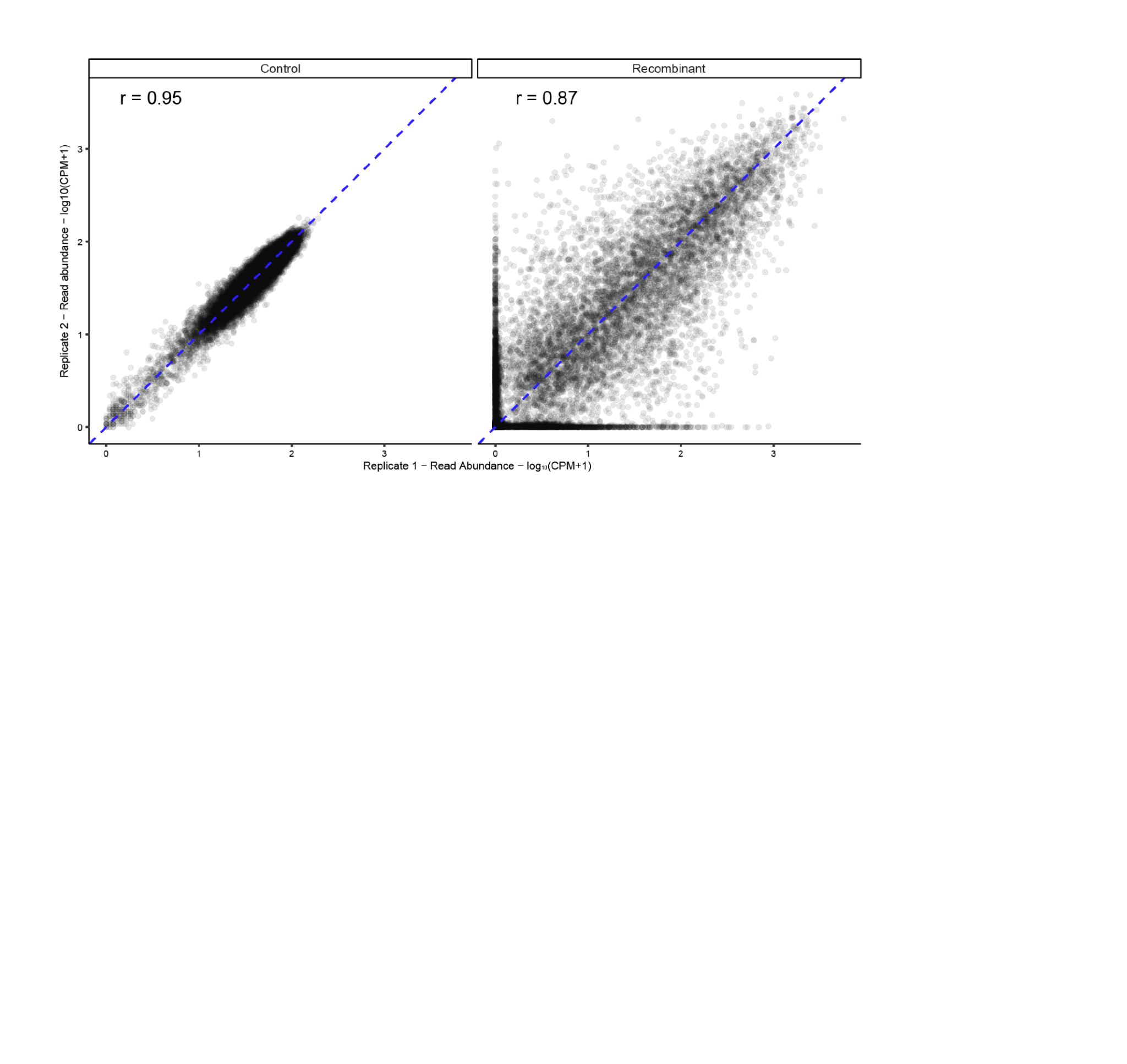

**Supplementary Data 4: Replicate correlation, donor-binding loop and donor screen**

Read abundance of library members in target-binding loop and target screen. Results are shown for two replicates for the control, amplifying only the barcode region, and recombinant, amplifying the barcode region across the recombination junction. Identity line shown as blue dashed line. r; Pearson correlation coefficient.

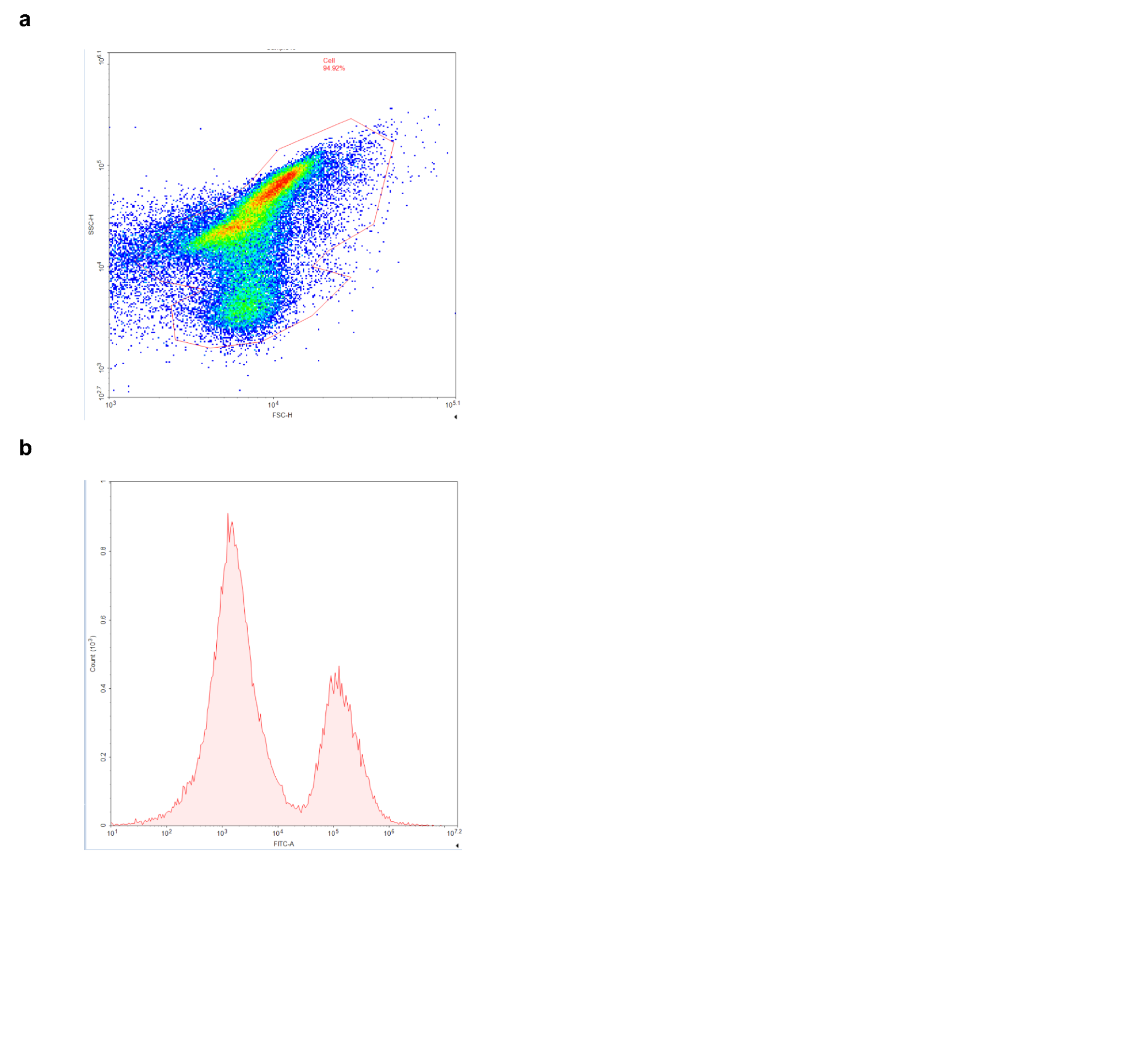

**Supplementary Data 5: Example of flow cytometry gating strategy**

(**a**) Gating on FSC/SSC of cells.

(**b**) Resulting histogram of FITC-A after gating in (a). The mean fluorescence intensity (MFI) of the lower plot was recorded for all experiments.

**Supplementary Note 1: Discussion of structural variants in genome insertion assay**

We developed a bioinformatics pipeline to detect possible structural variants in our genomic insertion assay that are independent from the donor plasmid (**Supplementary Table 3**). The most common structural variant (SV) appears to be a transposition event of the IS4 element located at NZ_CP053602.1:4413631-4415070 to NZ_CP053602.1:2604719, an intergenic sequence downstream of the outer membrane protein assembly factor *bamD* and upstream of the ribosome-associated inhibitor A *raiA*. This was detected at allele frequency (AF) > 0.5. Upon visual inspection of the alignments in IGV, it appears to be fixed at this position in several different experiments. Furthermore, no sequence similarity was detected between this SV and the bridge RNAs used in the respective experiments where the SV was detected. We therefore reasoned that this SV is unlikely to be due to the bridge RNA recombination mechanism.

A total of 17 structural variants were detected where one or both ends of the variant overlapped with a detected insertion site, with 10 of these replicating across >1 biological replicate. This is evidence that these structural variants were RNA-guided and independent of the donor plasmid. Overall, these variants were rare, with an average AF of 0.005 (range 0.001-0.023). We see one site in particular, located at NZ_CP053602.1:284995, acts as a recombination site for several different recombination events across different bridge RNA target-binding loops (all with the WT donor-binding loop), including 2 inversions and 6 deletions, that range in size from 50 kb to 537 kb. The genomic sequence at NZ_CP053602.1:284995 is ACGATATCTTG, which closely resembles the ACAGTATCTTG donor sequence (Levenshtein distance = 2). This suggests that such genomic rearrangements, if undesired, could be mitigated by more optimal donor-binding loop sequence design, analogous to CRISPR guide RNA design algorithms for choosing more specific spacer sequences.

Almost all structural variants are low-frequency mutations (AF < 0.05) in this experimental context, several with as little as 2 supporting reads. Due to their low frequency and large size, it is difficult to say with certainty if they represent insertions, deletions, or if they may be part of more complex genomic rearrangements. Intriguingly, one structural variant that is supported by 2 reads in 1 replicate suggests a ~1.13 Mb (NZ_CP053602.1:4106991-681788) deletion between two donor-like genomic sites, indicating the potential of the bridge mechanism to engineer long-range structural variants. Further analysis of bridge RNA-guided structural variants, both in their natural context and in the context of genome engineering, is an exciting direction for future research.
